# Ghost W chromosomes and unique genome architecture in ghost moths of the family Hepialidae

**DOI:** 10.1101/2023.09.03.556148

**Authors:** Anna Chung Voleníková, Ken Sahara, Jana Štundlová, Martina Dalíková, Petr Koutecký, Patrick Grof-Tisza, Thomas J. Simonsen, Michal Žurovec, Irena Provazníková, James R. Walters, František Marec, Petr Nguyen

**Author notes:** **Corresponding author:** Petr Nguyen, Faculty of Science, University of South Bohemia, Branišovská 1760, 37005 České Budějovice, Czech Republic.

## Abstract

The classical model of sex chromosome evolution has been recently challenged in moths and butterflies (Lepidoptera). According to the current hypothesis, the adoption of a supernumerary chromosome may have driven the transition from the Z0 to the WZ sex chromosome system in females. However, the evolutionary history of the W chromosome remains enigmatic, especially in the early-diverging lepidopteran lineages. In ghost moths of the family Hepialidae, one of the most basal lepidopteran clades, there is conflicting evidence regarding their sex chromosomes. In this study, we aimed to clarify the status of the hepialid W chromosome. Using cytogenetics and genomics, we investigated the karyotype, sex chromosomes, genome size and repeatome of multiple ghost moth species and reconstructed basic phylogenetic relationships in the group. Our data show that Hepialidae have unusually large genomes (reaching up to 1C = 3 Gb) and are the oldest known lepidopteran clade with a W chromosome. However, the W does not form a typical heterochromatin body in polyploid nuclei, known as sex chromatin, previously employed to detect the presence of W chromosomes across Lepidoptera. Moreover, in some species, the W does not exhibit distinct repeat content and can escape detection via methods that rely on W-specific sequences. Analysis of the Z chromosome confirmed highly conserved gene content, arguing for a possible origin of the hepialid W chromosome from a B chromosome. We hypothesize that the mechanism underlying the formation of sex chromatin could be used in future research to study the origin of the W chromosome.

## 1. INTRODUCTION

Genetic determination of sex by sex chromosomes has evolved independently many times in diverse taxa (**Bachtrog et al. 2014**). According to the classical model (**Charlesworth 1991**), differentiated sex chromosomes arise from a pair of autosomes in which one homolog acquires a sex-determining function. Subsequently, mutations favorable to the corresponding sex accumulate on the chromosome, and recombination is suppressed to tighten their linkage to the sex-determining region. Further expansion of the non-recombining region may eventually lead to gradual degeneration and genetic erosion of the sex-limited chromosome (Y or W), while its evolutionary counterpart (X or Z) retains its genes and autosome-like features (reviewed in **Charlesworth 2021**). The resulting difference in gene dose often needs to be compensated which can be done either by overexpressing or silencing the X- or Z-linked genes in the heterozygous or homozygous sex, respectively, to maintain ancestral expression levels (**Ohno 1967, Marín et al. 2000, Gu and Walters 2017**). The evolution of sex chromosomes follows this scenario in vertebrate lineages such as mammals (**Lahn and Page 1999**) and birds (**Handley et al. 2004**), but also in plants (**Filatov 2005**).

Recently, a growing body of evidence from non-model species has challenged the classical hypothesis of sex chromosome evolution and questioned its major predictions, leading to a paradigm shift in our understanding (**Furman et al. 2021, Johnson Pokorná and Reifová 2021, Kratochvíl et al. 2021**). According to the newly proposed hypotheses, the suppression of recombination and the resulting differentiation of sex chromosomes are not necessarily driven by sexually antagonistic selection but may be the consequence of neutral processes (**Jeffries et al. 2021, Perrin 2021**). The same outcome can arise by other means, such as epigenetic changes (**Zhang et al. 2008**), possibly resulting from the insertion of transposable elements (**Kent et al. 2017**). Epigenetic regulation, especially DNA methylation, may also play a key role in sex determination in systems with low sequence diversity of sex-determining genes (**Weber and Capel 2021, Zhang et al. 2022**). Sex chromosome turnover is common in groups with both homomorphic and heteromorphic sex chromosome systems (**Vicoso and Bachrog 2013, Jeffries et al. 2018**), and dosage compensation may occur on a gene-by-gene basis rather than at the whole chromosome level (reviewed in **Gu and Walters 2017**; **Fruchard et al. 2020**). Last but not least, horizontal gene transfer or supernumerary (B) chromosomes may provide genetic material for the *de novo* emergence of non-homologous sex chromosomes (**Fraïsse et al. 2017, Clark and Kocher 2019**).

Evolution of a sex chromosome from a B chromosome has recently been proposed in several animal taxa. B chromosomes are additional nuclear elements thought to have arisen by irregular rearrangements of the standard karyotype or interspecific hybridization. As a result, their genetic content, frequency and transmission rates are extremely variable (for a review, see **Johnson Pokorná and Reifová 2021**). Remarkably, they can undergo cellular domestication and take over the role of a sex chromosome. This can be achieved by acquisition of a sex-determining gene (**Clark and Kocher 2019**), fusion with an existing sex chromosome (**Conte et al. 2021**) or pairing with an X or Z univalent in X0/XX (♂/♀) or Z0/ZZ (♀/♂) systems (**Nokkala et al. 2003**).

Moths and butterflies (Lepidoptera) represent a prominent taxon long hypothesized to have sex chromosomes of non-canonical origin. In the clade Ditrysia, comprising 96% of lepidopteran species (**Van Nieukerken et al. 2011**), sex is typically determined by the WZ/ZZ (♀/♂) sex chromosome system with a gene-poor, heterochromatic W chromosome in females (**Traut et al. 2007**). However, the earliest-diverged lepidopteran lineages and their sister order, the caddisflies (Trichoptera), lack this element and possess Z0/ZZ (♀/♂) sex chromosomes (**Traut and Marec 1997, Marec and Novák 1998**). One long-standing hypothesis to explain this pattern is that the lepidopteran W chromosome evolved from the ancestral Z0 constitution via a Z-autosome fusion, followed by a rapid degeneration of the maternally inherited homolog that gave rise to the W chromosome (reviewed in **Sahara et al. 2012**). This scenario seemed plausible because sex chromosome-autosome fusions are indeed very common in Lepidoptera (**Nguyen et al. 2013, Nguyen and Carabajal Paladino 2016, Mongue et al. 2017, Carabajal Paladino et al. 2019, Yoshido et al. 2020**).

However, **Dalíková et al. (2017)** and **Fraïsse et al. (2017)** reported a conserved gene content of the Z chromosome in both non-ditrysian and ditrysian Lepidoptera, which does not support the fusion origin of the ditrysian W chromosome. It has been proposed that the W chromosome has a non-canonical origin and may have evolved by recruitment of a B chromosome (see discussion in **Fraïsse et al. 2017**). Therefore, moths and butterflies present a promising lineage in which to investigate the non-canonical evolution of sex chromosomes (**Furman et al. 2021**).

One of the crucial - and controversial - details yet to be resolved is when the W chromosome evolved in the Lepidoptera. **Traut and Marec (1996)** examined numerous lepidopteran species for the presence of sex chromatin, i.e. a female-specific heterochromatin body, which is formed by thousands of copies of the W chromosome in polyploid tissues. This distinct cytological feature has routinely been used as a proxy for the presence of the W chromosome, especially in species for which chromosome preparations were difficult to obtain due to specific larval biology or specimen size, such as non-ditrysian micromoths. Distribution of sex chromatin placed the W origin at the base of Ditrysia. Using the same line of evidence, **Lukhtanov (2000)** proposed that the W chromosome emerged in the common ancestor of Ditrysia and its closest outgroup, the Tischeridae (Fig. 1). However, the presence of a putative W chromosome was previously reported in the female karyotype of *Endoclita sinensis*, a representative of the much earlier-diverging clade Hepialidae (**Kawazoe 1987**), which would contradict the above two hypotheses. Interestingly, two other hepialid species tested negative for sex-chromatin, suggesting absence of the W in their genomes (**Traut and Marec 1996, Lukhtanov 2000**).

**Fig. 1.**
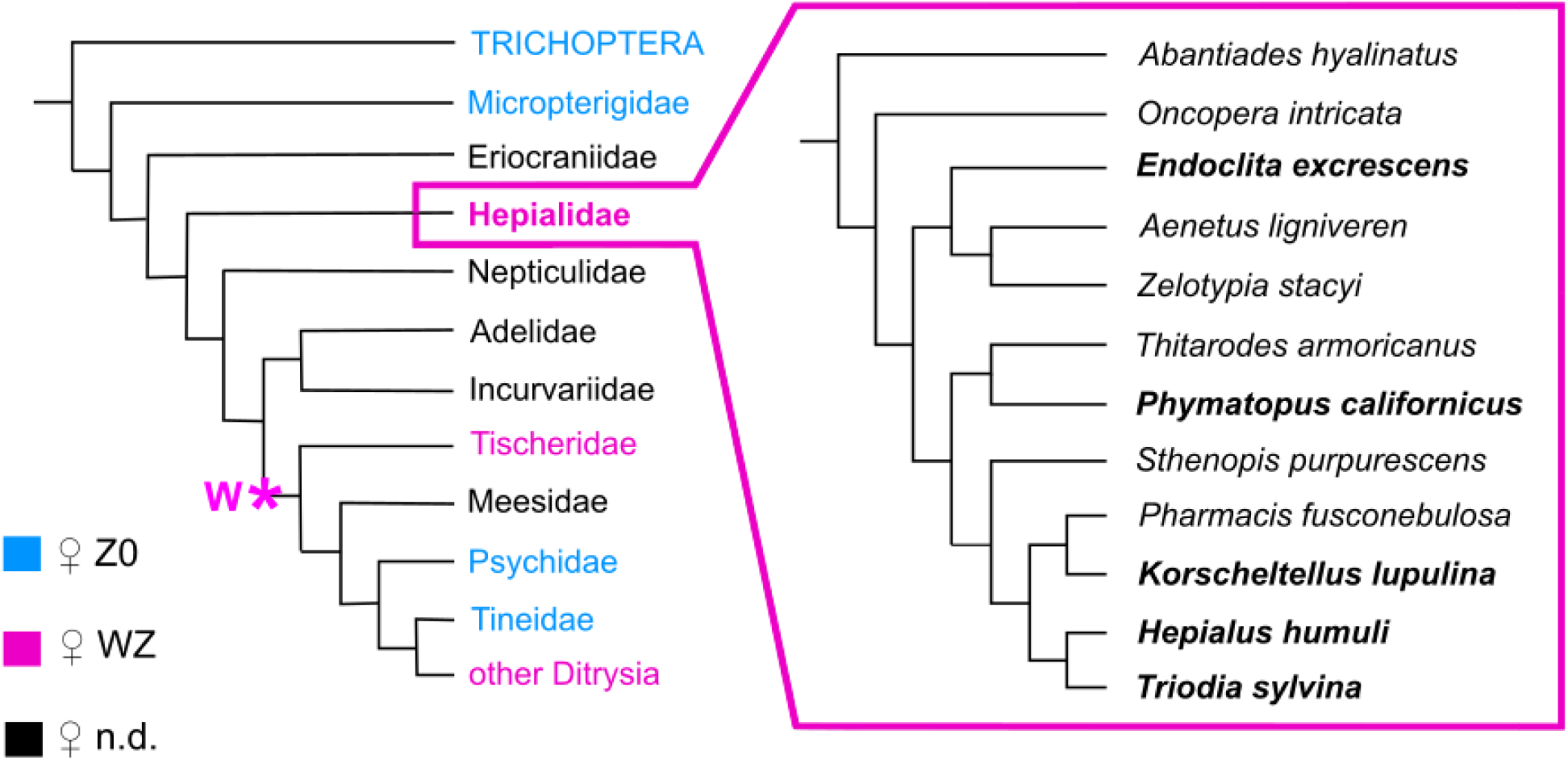
Lepidopteran phylogeny and reconstruction of phylogenetic relationships in Hepialidae. Lepidopteran phylogeny based on **Mitter et al. (2017)**. Clades with ♀Z0/♂ZZ sex chromosome system are in blue, ♀WZ/♂ZZ in magenta; n.d. - sex chromosomes not determined. Cytogenetic data on sex chromosome systems were obtained from published studies (Kawazoe 1987, Sahara et al. 2012; Dalíková et al. 2017; Hejníčková et al. 2019). Putative origin of W chromosome is indicated by asterisk. Hepialidae species examined in this study (in bold) were found in three separate branches, suggesting that the sampling represents the majority of Hepialidae.

In this work, we aimed to investigate the W chromosome origin in Lepidoptera by examining the sex chromosomes of enigmatic ghost moths of the family Hepialidae. By combining classical and molecular cytogenetics, genomics, and phylogenetics, we provide a new perspective on the evolution of sex chromosomes in Lepidoptera and uncover unique features of Hepialidae genomes.

## 2. RESULTS

### 2.1 Phylogeny of Hepialidae

To provide evolutionary context for our data, we reconstructed the phylogenetic relationships of the five studied hepialids and an additional seven ghost moth species. All phylogenetic analyzes revealed a very similar topology (for individual trees with maximum likelihood (ML) and Bayesian inference (BI) see Figs S1 and S2, respectively). Phylogenetic relationships among hepialid species were for the most part strongly supported and stable, including the phylogenetic positions of focal species, i.e., *Endoclita excrescens*, *Hepialus humuli*, *Korscheltellus lupulina*, *Phymatopus californicus*, and *Triodia sylvina*, in both ML and BI analyses. The species of interest did not form a distinct clade within the family and were scattered across the phylogenetic tree (Fig. 1) as follows: (i) *E. excrescens* proved to be a sister taxon to *Aenetus ligniveren* + *Zelotypia stacyi*; (ii) *P. californicus* proved to be closely related to *Thitarodes armoricanus*, together forming a sister clade to (iii) the clade with other five hepialid species, including *H. humuli*, *K. lupulina*, and *T. sylvina*.

### 2.2 Sex chromatin and karyotype analysis

Assays for sex-specific heterochromatin were performed in three hepialid species, namely *E. excrescens*, *P. californicus*, and *H. humuli*. No conspicuous heterochromatin body was detected in the polyploid tissues of the Malpighian tubules in either sex of tested species (Fig. 2a–d; Fig. S3). Heterochromatin grains were observed in some nuclei in *P. californicus* (Fig. S3c). However, their distribution and size varied and they were present in both sexes, which suggests their autosomal origin.

**Fig. 2.**
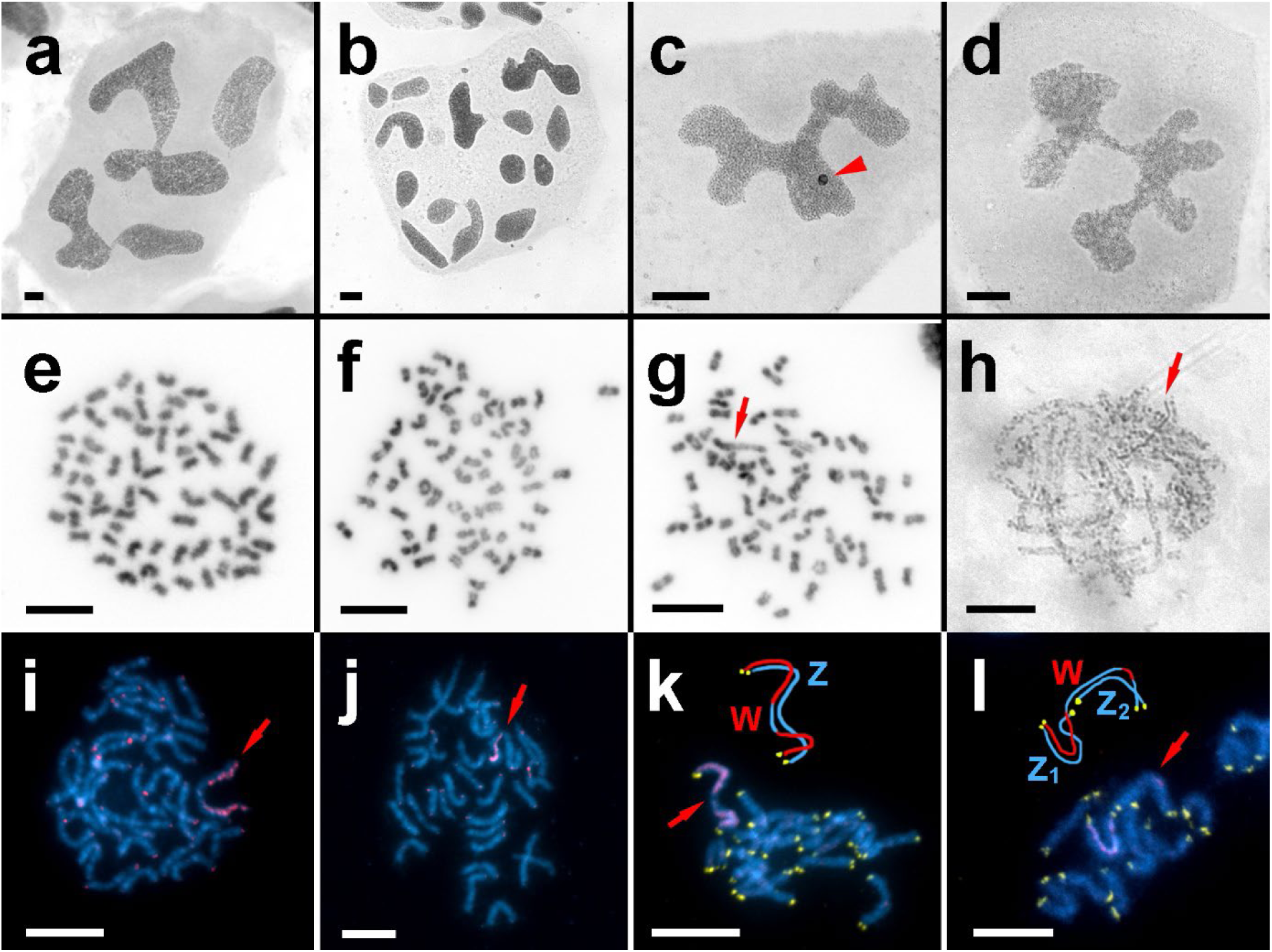
Sex chromatin assay, karyotypes, and W chromosome detection in Hepialidae. **(a–d)** Sex chromatin assay. No heteropycnotic body was present in the polyploid nuclei of the Malpighian tubules of females **(a)** and males **(b)** of *Hepialus humuli*. For comparison, conspicuous sex chromatin (arrowhead) was visible in females of the Mediterranean flour moth, *Ephestia kuehniella* (Pyralidae) **(c)** but not in males **(d)**. Bar **(a–d)** = 20 µm. **(e–h)** Mitotic and meiotic chromosomes stained with orcein. The ancestral hepialid karyotype consisting of 2n = 64 chromosomes represented by female **(e)** and male **(f)** mitotic nuclei of *Phymatopus californicus*. An alternative chromosomal race with a large neo-W chromosome (arrow) was detected in a subset of *P. californicus* females **(g)**. Deeply stained heterochromatic W chromosome (arrow) in a pachytene oocyte of *Korscheltellus lupulina* **(h)**. Bar **(e– h)** = 10 µm. **(i–l)** Detection of the W chromosome in female pachytene nuclei. Genomic *in situ* hybridization (GISH) confirmed the presence of the W chromosome in ancestral **(i)** and neo-W chromosome **(j)** races of *P. californicus*, showing distinct patterns of W differentiation (red). Female gDNA probe in red, DAPI staining in blue, position of W chromosome indicated by arrow. GISH combined with telomeric probe (yellow) identified WZ bivalent and WZ_1_Z_2_ trivalent in the ancestral **(k)** and neo-W chromosome **(l)** races, respectively (indicated by arrows and schematized above nuclei). Bar **(i–l)** = 10 µm.

Mitotic and meiotic complements from gonadal or brain tissue were examined in all examined species and their chromosome number was determined to be n = 32 in both sexes (Fig. 2e,f; Fig. S4a–h). The presence of 1–2 supernumerary (B) chromosomes was detected in one male and one female specimen of *H. humuli* from the same population. These chromosomes were smaller than the rest of the complement and differed in shape (Fig. S4e). In addition, distinct constriction-like morphologies were observed on mitotic chromosomes at the (pro)metaphase stage (Fig. 2e–g; Fig. S4a,d,e,h), which is unusual for lepidopteran holocentric chromosomes. However, early anaphase nuclei showed parallel separation of chromatids (Fig. S4b,g), typical for holocentric chromosomes.

The absence of sex chromatin has previously been interpreted as the absence of a W chromosome (**Traut and Marec 1996**). However, the even number of chromosomes observed in both sexes contradicts this and suggests that WZ/ZZ (♀/♂) is the likely sex chromosome constitution in Hepialidae. Indeed, a heterochromatinized W chromosome visible as a deeply stained DAPI-positive thread was found in female pachytene nuclei of *K. lupulina* (Fig. 2h), *P. californicus* (not shown), and *T. sylvina* (Fig. S4i). Remarkably, the W chromosome was polymorphic in females of *P. californicus,* dividing them into two groups. In the first group, females had 2n = 64 chromosomes and a W chromosome whose size was comparable to autosomes (Fig. 2f). In the second group, females had only 2n = 63 chromosomes and a large W element was present in their complement, indicating the possible presence of a neo-W chromosome (Fig. 2g). In *H. humuli*, obtained female pachytene nuclei were not sufficient to directly assess the status of the W chromosome.

### 2.3 Detection of the W chromosome

To confirm the karyotyping results, we visualized the W chromosome using genomic *in situ* hybridization (GISH) by hybridizing a fluorescently labeled female genomic DNA (gDNA) probe on female nuclei, together with excess of unlabeled competitor male gDNA. The male competitor gDNA hybridizes to sequences common to both sexes, leaving only female-enriched or female-specific sequences, which highlight the W chromosome. Indeed, a differentiated W chromosome was observed in *E. excrescens*, *K. lupulina* and *P. californicus* (Fig. 2i–l; Fig. S5a–f).

GISH also clearly demonstrated the existence of two chromosomal races in *P. californicus* females, where the female-derived genomic gDNA probe labeled either the whole W chromosome except for a short central region (Fig. 2i) or only half of the extraordinarily large W chromosome (Fig. 2j). GISH combined with telomeric fluorescence *in situ* hybridization (telo-FISH) suggests that the first case probably represents an ancestral W chromosome paired with a single Z chromosome (Fig. 2k) seen also in the other hepialids. In contrast, in the second race, the W chromosome probably fused with an autosome, resulting in a large neo-W chromosome, which consisted of a heterochromatic part corresponding to the ancestral W chromosome and a neo part of autosomal origin with only a small differentiated region (Fig. 2j). Accordingly, two Z chromosomes paired with the neo-W during meiosis represent the ancestral Z designated as Z_1_ and Z_2_ of autosomal origin. These chromosomal races were also distinguished as separated haplotypes by three single nucleotide polymorphisms in the *cytochrome c oxidase subunit I* gene (<u>Supplementary data 1;</u> accession numbers OR083623, OR083624). In our sampling (N=56 females), the neo-W chromosome-linked haplotype was more frequent (66%) than the haplotype associated with the ancestral sex chromosome constitution (34%).

Surprisingly, GISH failed to identify the W chromosome in *T. sylvina* and *H. humuli*, as no chromosome was labeled by the female gDNA probe. To further investigate the sex-specific/enriched segments of the female genome, comparative genomic hybridization (CGH) was performed, in which identical amounts of both female and male fluorescently labeled gDNA probes were hybridized competitively onto female chromosomes. No sex-specific differences were detected and the gDNA probes labeled all chromosomes equally. This was particularly noteworthy in the case of *T. sylvina*, where the highly heterochromatinized W chromosome was otherwise distinguishable by the DAPI-positive staining pattern (Fig. S5g–j). Because of the lack of material in *H. humuli*, we could not examine the pachytene nuclei in females of this species, but no major female-specific/enriched hybridization signals were observed on mitotic or interphase nuclei (Fig. S5k-n).

### 2.4 Genome size measurements

As GISH and CGH failed to provide direct evidence for presence of W chromosome in *H. humuli* and *T. sylvina*, we tried to further corroborate the presence of the W chromosome by comparing genome sizes between sexes by means of flow cytometry. If the W chromosome is absent in female karyotype, female genome should be smaller on average than male genome (**Hejníčková et al. 2019**), otherwise there should not be a significant difference in genome size between sexes. In addition to *H. humuli* and *T. sylvina*, the hepialid *P. californicus* was tested as a control species with a cytogenetically confirmed WZ/ZZ (♀/♂) sex chromosome system. At least five measurements of sufficient quality were obtained for each species and sex.

The genome size of *H. humuli* was determined to be 1C = 2.65 ± 0.06 pg (mean ± standard error) for females and 1C = 2.68 ± 0.07 pg for males, corresponding to a mean 1C = 2.61 Gb for this species. In *T. sylvina*, female genome size was 1C = 3.15 ± 0.05 pg and male 1C = 3.14 ± 0.03 pg with a mean of 1C = 3.08 Gb. In *P. californicus*, genome size was 1C = 1.62 ± 0.03 pg in females and 1.62 ± 0.02 pg in males with a mean of 1C = 1.58 Gb. In all cases, the difference in genome sizes between sexes was not statistically significant (*P* > 0.05; Table 1), which indicates the presence of the W chromosome in all tested species. Examples of flow cytometric histograms are shown in Fig. S6.

**Table 1.** Hepialidae genome sizes and comparison between the sexes^a^.

| Species | Sex | Control species | n | Mean ± SEM<sup>b</sup><br>(pg) | tvalue | Pvalue | F-test to<br>compare<br>variances | Average<br>(Gb) |
| --- | --- | --- | --- | --- | --- | --- | --- | --- |
|  |  | 1C (pg) |  |  |  |  |  |  |
| *Triodia*<br>*sylvina* | M | *Pyrrhocoris*<br>*apterus*<br>1C = 1.34 | 6 | 3.143 ± 0.031 | 0.429 | 0.678 | *P* > 0.05 | 3.08 |
|  | F |  | 5 | 3.153 ± 0.047 |  |  |  |  |
| *Hepialus*<br>*humuli* | M |  | 6 | 2.678 ± 0.069 | 0.745 | 0.473 | *P* > 0.05 | 2.61 |
|  | F |  | 6 | 2.651 ± 0.055 |  |  |  |  |
| *Phymatopus*<br>*californicus* | M | *Ephestia*<br>*kuehniella*<br>1 C = 0.45 | 9 | 1.619 ± 0.018 | 0.459 | 0.654 | *P* > 0.05 | 1.58 |
|  | F |  | 7 | 1.622 ± 0.034 |  |  |  |  |

<sup>a</sup> Statistically analyzed with unpaired two-tailed *t*-test for equal variances using GraphPad Prism 6 (GraphPad Software Inc., San Diego, CA, USA)
<sup>b</sup> SEM, standard error of the mean

### 2.5 Analysis of repetitive elements

GISH and CGH successfully detected the W chromosome in *P. californicus* but failed in *H. humuli* and *T. sylvina* (see section 2.3). We hypothesized that this is due to the repeat landscape of the W chromosome. Based on the percentage of clustered reads in RepeatExplorer2 analysis, the repetitive component of genome was estimated to form about 51.5% in *P. californicus,* 57.2% in *H. humuli* and 68.6% in *T. sylvina*. Most of the classified repeats were transposable elements (TEs) mainly from the retrotransposon class with LINE elements being the most prominent group except in *T. sylvina* where LINE and LTR elements make up very similar genome proportions (Fig. 3a). The satellite DNA formed only a small fraction of the repeats and the genome proportion formed by this type of repeats ranged from 0.5% in *P. californicus* to 2.9% in *T. sylvina* (Fig. 3a). As RepeatExplorer2 pipeline omits microsatellites, we used Tandem Repeat Finder to study tandem repeats with monomers up to 50 bp. The results showed only two to four different micro and minisatellites with genome proportions higher than 0.01 % in each species and the highest estimated proportion of one monomer was 0.05% (Table S1). Interestingly, one of these satellites in *T. sylvina and P. californicus* was the insect telomeric repeat TTAGG*_n_* (Table S1), while in *H. humuli* the telomeric repeat forms only about 0.003%.

**Fig. 3.**
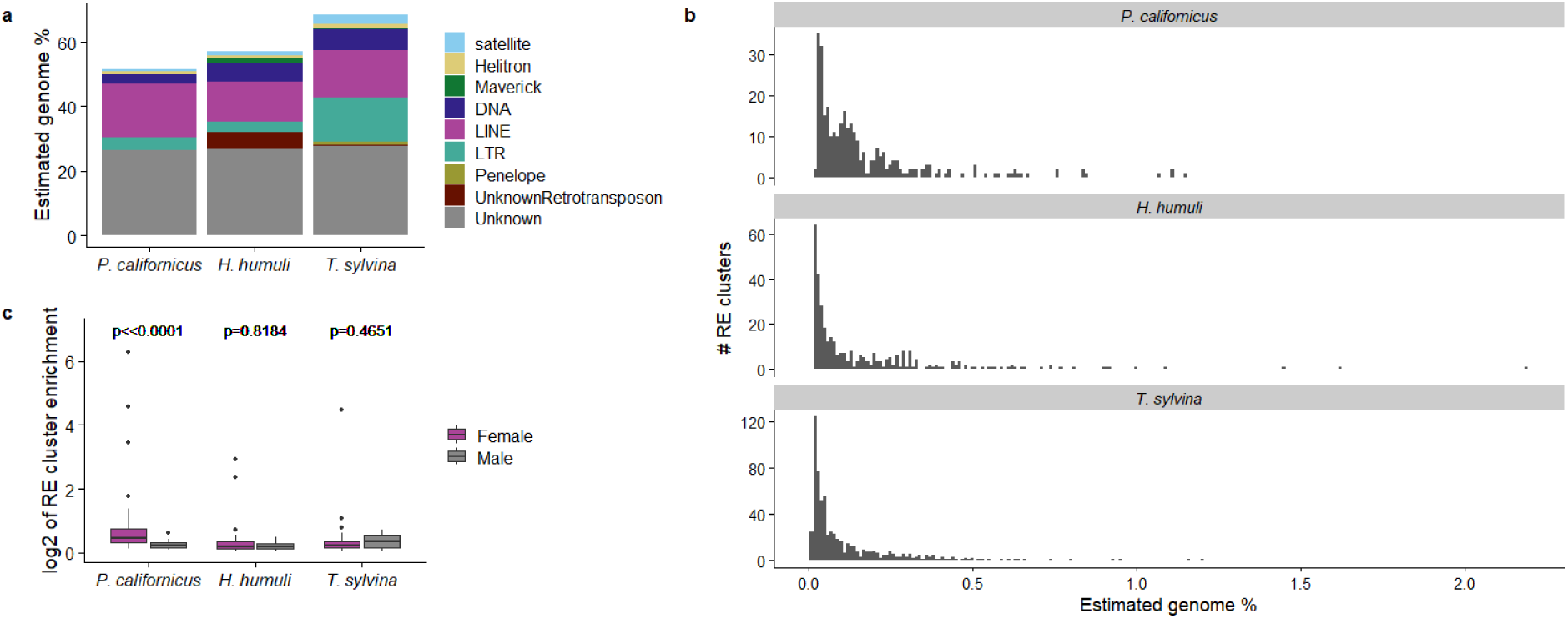
Results of RepeatExplorer2 analyses. **(a)** Genome proportion and classification of repeats identified by single male reads clustering. **(b)** Abundance of RE clusters in genomes of hepialid males. **(c)** Comparison of sex biased repeats between sexes of ghost moths species using Mann-Whitney-Wilcoxon test. In all cases Shapiro-Wilk normality test resulted in *P* < 0.05 for non-normal distribution of the data. Results show significant difference in male and female cluster enrichment in *Phymatopus californicus* (*P* = 3*10^−8^), but not in *Hepialus humuli* (*P* = 0.818) or *Triodia sylvina* (*P* = 0.465).

To analyze repeat accumulation on the W in these species, we performed RepeatExplorer2 comparative analysis between one male and one female sample with three independent read samplings. Each species exhibited several statistically significant (two sample *t*-test with unequal variances, *P* < 0.05) female enriched repeats, namely 39 clusters in *P. californicus*, 19 in *H. humuli* and 43 in *T. sylvina*, with the highest observed female enrichment 78x in *P. californicus* (Fig. 3c). However, in all cases there were also clusters exhibiting statistically significant male enrichment, which can be considered as intraspecific variability or technical bias. To compare the level of female and male enrichment we performed Mann-Whitney-Wilcoxon test, which showed that only *P. californicus* exhibits significantly higher female bias (Fig. 3c), suggesting that only in this species there is significant accumulation of W-enriched repeats.

### 2.6 Z-chromosome content in *Phymatopus californicus*

To determine the evolutionary origin of the W chromosome, comparison of the Z chromosome gene content between a hepialid representative, *P. californicus*, and a ditrysian model species, the silkworm *Bombyx mori* (Bombycidae), was performed. Array comparative genomic hybridization (aCGH) was used to test the linkage of 3,803 markers orthologous to a *B. mori* gene set. Of these, 148 orthologs were Z-linked and 3,452 were autosomal in *P. californicus*. Chromosomal assignment in the reference genome of *B. mori* revealed that 118 markers were Z-linked in both *B. mori* and *P. californicus* and were located along the entire Z chromosome of *B. mori* (Fig. 4). The remaining 30 orthologs that were Z-linked in *P. californicus* were found to be autosomal in *B. mori* and scattered over 17 chromosomes. Almost the same number, 28 orthologs, were autosomal in *P. californicus* but Z-linked and scatter along the Z chromosome in *B. mori*.

**Fig. 4.**
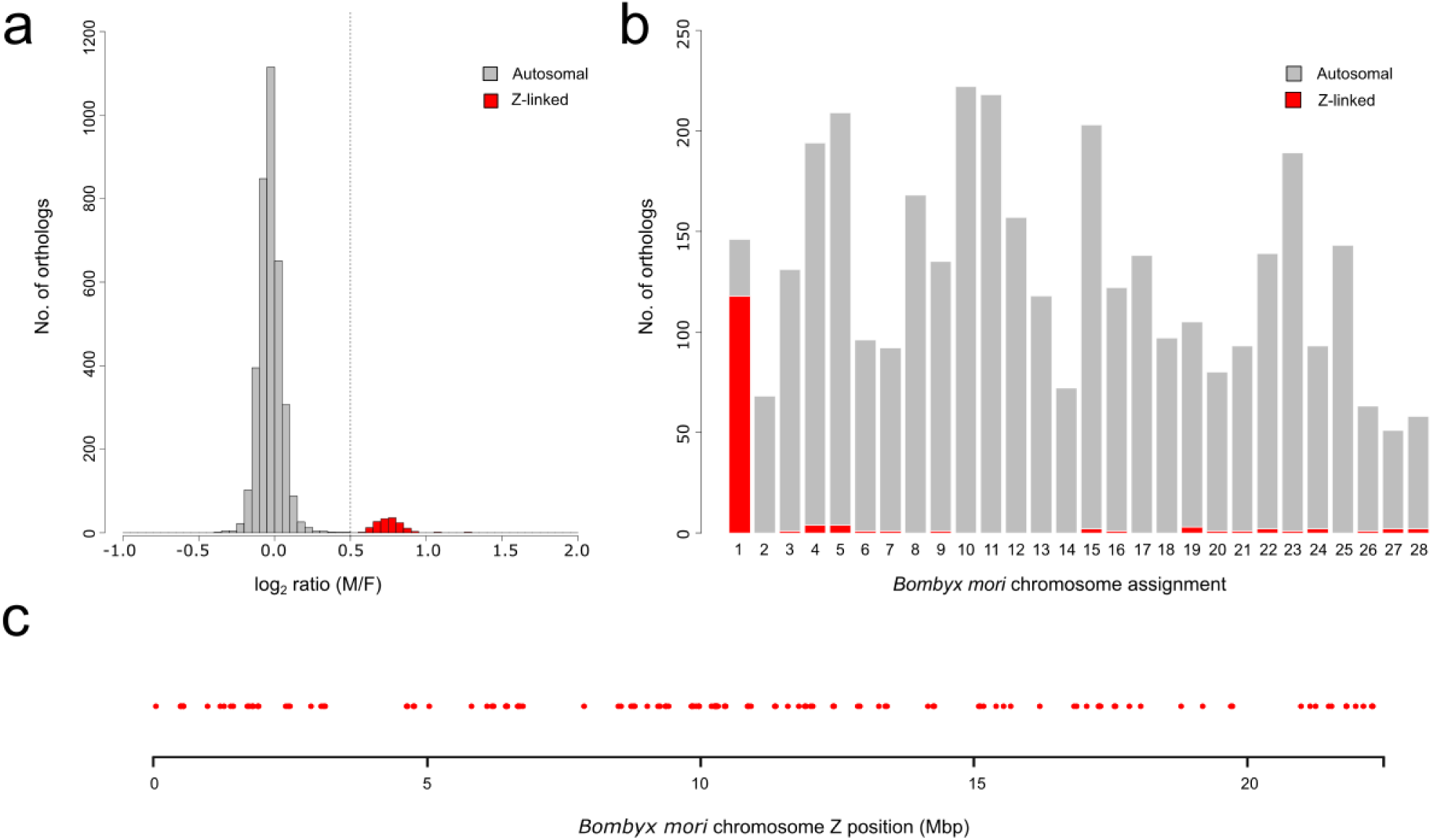
Array comparative genomic hybridization (aCGH) in *Phymatopus californicus*. **(a)** Bimodal distribution of log2 ratios of male-to-female signal intensities with a peak of putative autosomal orthologs (gray) at 0 and putative Z-linked orthologs (red) at approximately 0.75. **(b)** Assignment to the *Bombyx mori* reference genome shows that the majority of putative Z-linked orthologs in *P. californicus* (red) are also Z-linked in *B. mori* (assignment to chromosome 1). Autosomal orthologs are shown in gray. **(c)** Localization of orthologs that are Z-linked in both *P. californicus* and *B. mori* on the Z chromosome of *B. mori*.

## 3. DISCUSSION

To decipher the origin of the W chromosome in Lepidoptera and to clarify the conflicting evidence for the presence of the W chromosome in ghost moths (Hepialidae), we analyzed the karyotypes and sex chromosomes of five hepialid species. Their evolutionary relationships were unknown due to the lack of comprehensive phylogenetic hypothesis for Hepialidae. To date, the most extensive phylogeny has been based on morphological characters (**Grehan 2012**). A number of hepialids have been included in broader molecular phylogenetic studies (**Mutanen et al. 2010**, Wiegmann et al. 2012, **Regier et al. 2015**), but with little overlap in their datasets. To provide an evolutionary context for our results, we reconstructed the phylogenetic relationships of the five studied species plus species from seven additional hepialid genera. Both the maximum likelihood method (ML) and Bayesian inference (BI) yielded congruent topologies (Figs S1 and S2) with the examined ghost moths belonging to three lineages (Fig. 1), indicating that the sampling is representative for the family Hepialidae.

W-derived heterochromatin, also known as sex chromatin, has often been used as indirect evidence for the presence of the W chromosome in females of moths and butterflies (e.g. **Traut and Mosbacher 1968, Ennis 1976, Traut and Marec 1996**). Based on these data, it was hypothesized that the W chromosome is absent in early-diverging lepidopteran groups, including ghost moths. Confirming previous results in *Hepialus humuli* and *Triodia sylvina* (**Traut and Marec 1996, Lukhtanov 2000**), we found no heterochromatin bodies in females and males of these species, nor did we observe sex chromatin in *Phymatopus californicus* or *Endoclita excrescens* (Fig. 2 and Fig. S3). Under the previous assumption that the lepidopteran W chromosome will always form sex chromatin, this observation would suggest a Z0/ZZ (♀/♂) sex chromosome system in this species. However, equal numbers of chromosomes in both sexes and/or genomic *in situ* hybridization support the presence of the W chromosome in females of all species examined (Fig. 2, Figs S4 and S5), corroborating the previous report of the W chromosome in *Endoclita sinensis* (Kawazoe 1978).

Remarkably, the hepialid W chromosome does not appear to form the sex chromatin body typical of ditrysian moths and *Tischeria* (**Traut and Marec 1996, Dalíková et al. 2017**). Sex chromatin formation is impaired in ditrysian species with young neo-W chromosomes formed by translocation or fusion of an autosome with the W chromosome, likely as a consequence of active transcription from the neo-W chromosome (**Marec and Traut 1994, Vlašánek et al. 2017**). The decay of sex chromatin has also been observed in geometrid moths with an undifferentiated W chromosome (**Hejníčková et al. 2021**), suggesting that the absence of sex chromatin may be related to the genetic content of the W chromosome. However, this is not the case in the Hepialidae. Here, the W chromosome is well differentiated and heterochromatinized, as shown by DAPI and orcein staining (Fig. 2, Figs S4 and S5) and genomic *in situ* hybridization (see below). Therefore, we hypothesize that the lack of a sex chromatin body in Hepialidae may be due to absence of the machinery underlying sex chromatin formation in Tischeriidae and Ditrysia.

The sequence composition of the W chromosome is important for its successful detection by molecular cytogenetic methods, such as genomic *in situ* hybridization (GISH) and comparative genomic hybridization (CGH) (Fig. 2 and Fig. S5). In *T. sylvina*, we observed a large and conspicuous DAPI-positive W chromosome in the female pachytene nuclei (Fig. S5g). However, CGH revealed no difference in the hybridization patterns of the male and female gDNA probes in this species (Fig. S5g–j). Similar results were also observed in *H. humuli*, but not in the other hepialids examined, in which the W chromosome was readily detectable via GISH and CGH (Fig. 2, Fig. S5). To test whether our results could be explained by the repeat landscape of the W chromosome, we compared the male and female repeatomes of *P. californicus, H. humuli* and *T. sylvina*, using the RepeatExplorer2 pipeline. Indeed, we found a significant female bias in repeat content in *P. californicus* but not in *H. humuli* and *T. sylvina* (Fig. 3c), which agrees well with our hybridization results. Thus, the W chromosome in *P. californicus* is occupied by repeats significantly enriched or specific for the W and detectable by GISH. However, the repeat landscape does not differ between the W chromosome and the rest of the genome in *H. humuli* and *T. sylvina*, which hampers detection of W chromosome by means of molecular cytogenetics.

Because molecular cytogenetics failed to visualize the W chromosome in *H. humuli* and *T. sylvina*, we further corroborated the karyotyping data in these species by flow cytometry, using the hepialid *P. californicus* as a control with clear WZ/ZZ (♀/♂) sex chromosomes (section 2.3). As previously shown in bagworms (Psychidae), significant genome size differences between sexes are to be expected in taxa with the Z0/ZZ (♀/♂) sex chromosome system and chromosome counts similar to our study group (**Hejníčková et al. 2019**). No significant differences were found between the sexes in any of the studied species. But the mean genome sizes of *P. californicus*, *H. humuli*, and *T. sylvina* correspond to 1C = 1.58 Gb, 2.61 Gb, and 3.08 Gb, respectively (Table 1, Fig. S6). This indicates that hepialids have the largest lepidopteran genomes known to date, as haploid genome size rarely exceeds 1 Gb in moths and butterflies (**Calatayud et al. 2016, Cheng et al. 2016, Liu et al. 2020** and references therein, **Animal Genome Size Database 2022**).

To further investigate the causes of large genome sizes in the Hepialidae, we conducted a study of the repetitive genome fraction in *H. humuli*, *T. sylvina* and *P. californicus.* According to **Talla et al. (2017)**, genome size in Lepidoptera correlates with the amount of transposable elements in the genome. The proportion of repeats increased with genome size and accounted for approximately 45% of the genome in *P. californicus*, 50% in *H. humuli*, and 60% in *T. sylvina* (Fig. 3a). Surprisingly, these values are comparable to the 46.8% repeat content in the considerably smaller 0.5 Gb genome of the silkworm *B. mori* (**Mita et al. 2004, Kawamoto et al. 2019**). Instead of a massive amplification of a few specific repeats or classes of repeats (cf. **Mora et al. 2020**), a wide range of various repeats seems to be responsible for the genome expansion in Hepialidae. In all studied genomes, the most abundant repeat occupies only about 2% of the genome (Fig. 3b). This repeat diversity, as well as the relatively low proportion of reads clustering as repetitive DNA, could be explained by a trend recently observed in plants (**Novák et al. 2020a**). Analysis of plants with large genomes revealed an abundance of degraded repeats turned from highly repetitive sequences into unique or low-copy-number sequences through mutations and rearrangements.

It has been suggested that a high proportion of repeats may be associated with an increased rate of chromosomal rearrangements (**Miller and Capy 2004**, cf. **Hill et al. 2019**). This is especially true for lepidopterans, as they typically have holocentric chromosomes (**Wolf 1994**). However, the diploid chromosome number of 2n = 64 chromosomes is shared by all studied ghost moths, indicating a remarkable stability of karyotypes in Hepialidae. It is worth noting that we observed distinct constriction-like morphologies on mitotic chromosomes in all hepialids examined (Fig. 2, Fig. S4). According to **Wolf (1994)**, the kinetochore plate covers only about 30–70% of the lepidopteran chromosomes and the chromosomes should thus be considered only functionally holocentric. It is tempting to speculate that genome expansion, which has led to an increase in chromosome length, has resulted in the emergence of functionally monocentric chromosomes providing an evolutionary constraint on chromosome fusions and fissions.

Nevertheless, deviations from the standard hepialid karyotype were observed in *H. humuli* and *P. californicus*. In *H. humuli*, we noticed 1–2 supernumerary B-chromosomes (Fig. S4e), a phenomenon described so far in more than 30 lepidopterans (**B-chrom Database 2022**). More importantly, two chromosomal races were found in *P. californicus*, one with standard and presumably ancestral sex chromosome constitution and the other with a neo-sex chromosome (Fig. 2). GISH combined with the detection of telomeric repeats revealed the presence of a large neo-W chromosome, probably resulting from a fusion of the ancestral W chromosome with an autosome. Consequently, two Z chromosomes paired in meiosis with the neo-W chromosome to form the WZ_1_Z_2_ trivalent. The Z_1_ chromosome paired with the differentiated ancestral part of the neo-W chromosome while the Z_2_ chromosome paired with the neo-part of autosomal origin. Interestingly, the neo-W race predominated in the sampled population and was associated with a mitochondrial haplotype characterized by three single nucleotide polymorphisms in a sequence of the *cytochrome c oxidase subunit I* gene (<u>Supplementary data 1</u>), suggesting successful establishment of the neo-W chromosome in the population. Evolutionarily young sex chromosomes coexisting within a species as sex chromosome races provide excellent opportunities to gain much-needed insights into the early stages of sex chromosome evolution (**Furman et al. 2021**), as has been shown in grasshoppers (**Bugrov et al. 2004**) and plants (**Grabowska-Joachimiak et al. 2015**). In addition, the formation of reproductive isolation or phylogeographic patterns of genetic differentiation can be studied in hybrid zones (**Beaudry et al. 2019, Grzywacz et al. 2019, Smith et al. 2019**).

To identify the origin of the Z_2_ chromosome, we used array comparative genomic hybridization (aCGH), which has been successfully used previously to determine the origin of neo-sex chromosomes in flies and butterflies (**Baker and Wilkinson 2010, Yoshido et al. 2020**). However, the identified sex-linked orthologs were mostly assigned to the Z chromosome of the silkworm *B. mori* (Fig. 4). The results confirmed that the ancestral Z chromosome content is highly conserved in Lepidoptera (**Dalíková et al. 2017, Fraïsse et al. 2017**). Of the 130 markers identified as Z-linked in *P. californicus*, 91% were also Z-linked in *B. mori*. Previously, only 73.1% of the 78 Z-linked orthologs of *B. mori* were shared with *T. sylvina* (**Fraïsse et al. 2017**). Moreover, the positions of our markers cover the entire length of the *B. mori* Z chromosome, suggesting conserved synteny of the entire linkage group across Lepidoptera, as proposed by **Dalíková et al. (2017)** (Fig. 4). Thus, the content of the ancestral Z chromosome apparently rules out its evolutionary origin in sex chromosome-autosome fusion.

While the ancestral WZ sex chromosome pair is well differentiated (cf. **Mongue et al 2017, Palmer et al. 2019**), the aCGH results suggest that the W chromosome-autosome fusion occurred only recently, probably less than 1 million years ago given the achiasmatic meiosis of lepidopteran females (see discussions in **Mongue et al. 2017** and Yoshido et al. 2020). We hypothesize that a short differentiated region detected by GISH on the neo-part of the neo-W chromosome of *P. californicus* (Fig. 2l) may have arisen from an intrachromosomal translocation or an inversion with breakpoint between the ancestral part and the neo-part of the neo-W chromosome.

The origin of the W chromosome in lepidopteran phylogeny remains unclear. Although it was proposed it is absent in early diverging clades, caution is warranted because this information was inferred from the absence of sex chromatin only. As discussed above, the sex chromatin is not a good proxy for presence of W chromosome in hepialids. Considering the confirmed presence of W chromosome in Hepialidae, we can now envision two equally parsimonious hypotheses regarding W chromosome origins in Lepidoptera. Both scenarios include three evolutionary events: either a single W emergence in a common ancestor of the Hepialidae and Ditrysia clades and two W losses in Psychidae and Tineidae, or three independent W origins in Hepialidae, Tischeriidae and Ditrysia. Further karyotype investigations in the earliest-diverging lepidopteran lineages are needed as well as comparisons of Z-linked gene content between Hepialidae, Ditrysia and other groups with a Z0/ZZ sex chromosome system, such as the mandibulate moths of the family Micropterigidae and the caddisflies.

The main obstacle in deciphering the evolutionary origin of lepidopteran W chromosomes lies in their molecular composition. W chromosomes have been shown to be packed with repetitive sequences that differ substantially due to their rapid turnover even among closely related species (**Vítková et al. 2007, Traut et al. 2013, Zrzavá et al. 2018, Lewis et al. 2021, Berner et al. 2023**). W-linked protein-coding genes were reported in only a handful of species with no overlap (**Gotter et al. 1999, Van’t Hof et al. 2013, Nagaraju et al. 2014, Fujii et al. 2015**). The lack of homology between W and Z chromosomes is not informative, as it is expected in both scenarios of the W chromosome origin. Under an autosomal origin, all coding sequences confined to the newly formed W chromosome would rapidly degenerate due to achiasmatic meiosis of lepidopteran females (reviewed in **Satomura et al. 2019**) and thus a complete absence of recombination. Regarding the origin of the W chromosome from a B chromosome, the B chromosomes generally arise from repetitive parts of the genome and continue to accumulate repeats and other genetic material (**Johnson Pokorná and Reifová 2021**).

Our results point to yet another line of evidence that may shed light on the origin of the W chromosome in Lepidoptera. We argue that the machinery underlying the silencing of W-linked repeats, including the formation of sex chromatin, can be used to investigate the origin of W chromosomes in Lepidoptera. Although no data are available on the mechanistic basis of sex chromatin formation in Lepidoptera, several models have been proposed in other taxa. In humans (**Hall and Lawrence 2010**), mice (**Maison et al. 2002**), and *Tetrahymena* (**Kataoka and Mochizuki 2015**), the interplay between non-coding RNAs and heterochromatin-associated proteins is thought to serve as a platform for the formation of highly organized heterochromatin. The formation of heterochromatin domains can be driven by liquid phase separation in *Drosophila* (**Strom et al. 2017**), while protein-protein interactions are thought to play a role in clustering heterochromatin from different chromosomes into a subnuclear compartment in budding yeast (**Ruault et al. 2021**). Similar to dosage compensation, epigenetic silencing is likely associated with early differentiation of sex chromosomes, i.e. degeneration of the W chromosome through the accumulation of repeats and pseudogenization of W-linked alleles. Thus, the mechanistic basis of these processes may allow us to test for a common origin of W chromosomes in hepialid and ditrysian moths despite the lack of sequence homology.

## 4. MATERIALS AND METHODS

### 4.1 Insects

Various life stages of ghost moths were collected in Japan (*Endoclita excrescens*), the Czech Republic (*Hepialus humuli*, *Triodia sylvina*, *Korscheltellus lupulina*), Austria (*Hepialus humuli*), and the USA (*Phymatopus californicus*). The exact localities are given in Table S2. In *E. excrescens*, larvae were reared with artificial diet (Insecta LFS, NOSAN corporation, Yokohama, Japan). For *H. humuli* and *P. californicus*, larvae were kept on the roots of *Daucus carota* until they reached the optimal stage for dissection. Hepialids collected as larvae were barcoded according to **Hebert et al. (2004)**. The Mediterranean flour moth *Ephestia kuehniella* (Lepidoptera, Pyralidae) and the firebug *Pyrrhocoris apterus* (Hemiptera, Pyrrhocoridae) used for genome size measurements were from laboratory strains kept at the Institute of Entomology BC CAS in České Budějovice (**Marec 1990, Socha et al. 2004**).

### 4.2 Molecular phylogeny

Sequences for the reconstruction of phylogenetic relationships were obtained either from transcriptomic data (*E. excrescens*, *H. humuli*, *P. californicus*, *T. sylvina, Thitarodes armoricanus, Dyseriocrania griseocapitella, Lophocorona astiptica*), genomic data (*K. lupulina*) or amplified by PCR from gDNA (*Abantiades hyalinatus*, *Oncopera intricata*, *Aenetus ligniveren*, *Zelotypia stacyi*, *Sthenopis purpurescens*, *Pharmacis fusconebulosa*). Details, accession numbers, and sequence extraction notes are provided in Table S3 and <u>Supplementary methods</u>. A dataset with 7 gene markers available in most species was generated for 12 hepialid species and 2 outgroups. Sequence alignment was performed independently for each gene using MAFFT v.7 (**Katoh et al. 2019**) with default settings. Individual alignments were visually inspected, checked for the presence of stop codons and concatenated in a single matrix of a total length of 4,057 bp in Geneious Prime v. 2021.1.1 (https://www.geneious.com). The alignment was deposited in the Dryad repository (https://doi.org/10.5061/dryad.w3r2280xb). The best partitioning scheme for the dataset and the best-fit model of molecular evolution for each subset were determined with PartitionFinder v. 2.1.1 (**Lanfear et al. 2016**) using the corrected Akaike information criterion (AICc). Phylogenetic relationships were reconstructed with maximum likelihood (ML) and Bayesian inference (BI) methods. The ML analysis was ran using RAXML v. 8.2.12 (**Stamatakis 2014**) with a separate GTRGAMMA model applied to each partition (Table S4). The best-scoring ML tree was selected from 100 independent iterations, each starting from a distinct randomized maximum parsimony tree, and the branch support values were calculated from 1000 bootstrap replicates. The BI analysis was conducted in MrBayes v. 3.2.7 (**Ronquist et al. 2012**). Models of molecular evolution were assigned to each partition, following the results of PartitionFinder analysis (Table S5). Two independent runs of 40 millions generations and four chains each sampled every 1000 generations were run simultaneously. Convergence and chain mixing of the runs were assessed by the mean standard deviation of split frequencies (< 0.01) and proper sampling of parameters (ESS values) summarized in Tracer v. 1.7.1 (**Rambaut et al. 2018**). The first 25% of sampled trees were discarded as a burn-in. Trees were visualized and edited using FigTree v. 1.4.3 (**Rambaut 2018**), Affinity Designer v. 1.10.4 (Serif, West Bridgford, UK) and Inkscape v. 1.2 (https://inkscape.org/).

### 4.3 Sex chromatin assay

Analysis of sex-specific heterochromatin was done based on Traut and Mosbacher, 1986. Briefly, Malpighian tubules were dissected from larvae in physiological solution, fixed for 2 min with Carnoy fixative (ethanol : chloroform : acetic acid, 6: 3: 1), and stained for 3 min with 1.5% lactic acetic orcein.

### 4.4 Chromosome preparations

Chromosome preparations for karyotyping and *in situ* hybridization were obtained from gonads and brains of late instar larvae (*E. excrescens, H. humuli, K. lupulina, P. californicus*) or pupae (*T. sylvina*) using previously described spreading techniques (**Mediouni et al. 2004**; **Yoshido et al. 2015** for *E. excrescens*). For chromosome counting, preparations were stained with 0.5 µg/ mL DAPI (4′,6-diamidino-2-phenylindole) in DABCO (1,4-diazabicyclo(2.2.2)-octane) antifade. Due to DAPI artifacts in *H. humuli* and *T. sylvina*, staining with 2% lactic acetic orcein for 1 hour at RT was used instead.

### 4.5 Genomic *in situ* hybridization (GISH)

Genomic DNA (gDNA) was isolated from males and females separately by standard phenol-chloroform extraction or cetyltrimethylammonium bromide as described in **Ferguson et al. (2021)**. Male gDNA was fragmented by boiling at 99°C for 10 min, and female gDNA was labeled with Cy3-dUTP using nick translation as described in **Hejníčková et al. (2019)**. Genomic *in situ* hybridization was carried out according to **Fuková et al. (2005)**.

### 4.6 Microscopy and image processing

The results were documented using Zeiss Axioplan 2 microscope (Carl Zeiss, Jena, Germany), Olympus CCD XM10 monochrome camera (Olympus Europa Holding, Hamburg, Germany) and Leica DM6000B fluorescence microscope (Leica Microsystems, Wetzlar, Germany). Images were captured with cellSens 1.9 imaging software (Olympus) and DFC350FX camera (Leica Microsystems) and processed using Adobe Photoshop Elements 2020 (Adobe Systems, San Jose, CA, USA).

### 4.7 Genome size measurements

Genome size for both sexes of *H. humuli*, *T. sylvina* and *P. californicus* was determined using flow cytometry. We used the firebug *Pyrrhocoris apterus* males and the Mediterranean flour moth *Ephestia kuehniella* (Pyralidae) males as internal standards. The genome size of *E. kuehniella* was previously measured as 1C = 0.45 pg (**Buntrock et al. 2012**). For *P. apterus* males, we used a value 1C = 1.34 pg, which is based on our own calibration (the mean fluorescence ratio of *P. apterus* male / *E. kuehniella* male averaged over 10 replicates was 2.969). All resulting values are therefore compatible with the published *E. kuehniella* genome size; note that previously published genome size of *P. apterus* based on less reliable method of Feulgen densitometry is slightly lower, 1C = 1.22 pg (**Bier and Müller 1969**). The heads of the standard and the measured specimen were macerated in nuclei isolation buffer and the resulting suspension was filtered and stained with propidium iodide, according to **Hejníčková et al. (2019)**. Samples were analyzed using Sysmex CyFlow Space flow cytometer (Sysmex Partec, Münster, Germany) equipped with a 100 mW 532 nm (green) solid-state laser and results were recorded and processed in FlowJo 10 software (TreeStar, Inc., Ashland, OR, USA). Genome size was calculated as the ratio of the mean fluorescence of the sample to the standard multiplied with the standard genome size. For *P. californicus*, we used *E. kuehniella* as the internal standard while for *H. humuli* and *T. sylvina* we used *P. apterus* due to large genome size difference exceeding the recommended 4-fold threshold to assure the measurement linearity (**Sliwinska et al. 2022**). To compare both sexes of *E. kuehniella*, males and females were measured against *P. apterus* males. Genome size differences of males and females were statistically analyzed with an unpaired two-tailed *t*-test for equal variances using GraphPad Prism 6 (GraphPad Software Inc., San Diego, CA, USA).

### 4.8 Repetitive DNA analysis

Repetitive genome content was examined in *H. humuli*, *P. californicus*, and *T. sylvina*. Whole-genome sequencing data were either obtained from NCBI (*T. sylvina*; accession numbers SRR5884531 and SRR5626447) or sequenced *de novo*. High-quality gDNA was isolated from male and female larvae of *H. humuli* and *P. californicus* using the MagAttract HMW DNA kit (Qiagen, Hilden, Germany) and sequenced at the Illumina platform by Novogene (HK) Co., Ltd. (Hong Kong, China). Raw reads were trimmed to a uniform length of 110 bp and filtered for average quality score at least 18 using Trimmomatic (v0.36, **Bolger et al. 2014**) and further preprocessed using the RepeatExplorer2 toolkit (**Novák et al. 2017**) according to **Novák et al. 2020b**. Repeats in each species were identified and annotated using Tandem Repeats Finder (**Benson 1999**), RepeatModeler (**Smit and Hubley, 2008-2015**) and RepeatExplorer2 pipelines (**Novák et al. 2013, Novák et al. 2020b**). To obtain an overview of repetitive fraction of genome in each species one male dataset subsampled to ca 0.3x genome coverage was analyzed by RepeatExplorer2 pipelines version 0.3.8.1 (**Novák et al. 2013, Novák et al. 2020b**). Repeats identified by this pipeline were annotated via RepeatExplorer2 automatic annotation with protein database metazoan v3 and using RepeatMasker (**Smit and Hubley, 2008-2015**). In case of a conflict in TE classification between these two approaches, the annotation with the longest matching region was considered if the difference was at least three times or the classification was downgraded to the level with consistent results. In the case of repeats annotated both as satellites and TEs, the tandem repeat classification was retained. As this tool omits short tandem repeats (**Novák et al. 2020b**), these were analyzed on the same subset of reads by Tandem Repeat Finder version 4.09 (**Benson 1999**) and Tandem Repeat Analysis Program (**Sobreira et al., 2006**). In both RepeatExplorer2 and Tandem Repeat Finder analysis, only repeats corresponding to at least 0.01 % of input data were considered. RepeatExplorer2 was also used for the analysis of sex-specific and enriched repeats, using 3,000,000 reads per species and three technical replicas per each sex. The resulting data were processed in R (version 4.2.1) and R Studio (**RStudio Team 2019, R Core Team 2020**).

### 4.9 Array comparative genomic hybridization (aCGH)

To determine Z chromosome content in *P. californicus*, array comparative genomic hybridization was performed according to **Baker and Wilkinson (2010)**. First, the transcriptome of *P. californicus* was assembled using available RNA-seq data (accession number SRR1021622). The raw reads were visualized using FastQC (**Andrews 2010**) and trimmed and quality filtered using Trimmomatic (v0.30, arguments ‘ILLUMINACLIP:${adapters.fa}:2:30:10:8:true LEADING:5 TRAILING:5 SLIDINGWINDOW:4:20’; **Bolger et al. 2014**). The resulting paired reads were then assembled by SOAPdenovo-trans-127mer with multiple odd k-mer sizes ranging from 21 to 81 (**Xie et al. 2014**) and Trinity (**Grabherr et al. 2011**). The resulting assemblies were concetenated and redundancy removed by the EvidentialGene pipeline (**Gilbert 2013**). The non-redundant transcriptome sequence was searched for orthologs of *B. mori* using HaMStR (**Ebersberger et al. 2009**). For each ortholog from the dataset, 60-mer oligonucleotide probes were designed in Agilent Technologies eArray Design Wizard (available online at https://earray.chem.agilent.com/earray/) and printed onto an Agilent-made custom microarray slide. Genomic DNA from males and females of *P. californicus* (neo-W chromosome race) was extracted separately using the MagAttract HMW DNA kit (Qiagen) and labeled using the SureTag Complete DNA Labeling Kit (Agilent) according to the protocol for Agilent Oligonucleotide Array-based CGH for Genomic DNA Analysis; the same protocol was used for hybridization with the Oligo aCGH/ChIP-on-chip Hybridization Kit (Agilent). Results were processed in Python (v. 3.4.1; https://python.org) using a custom script from **Yoshido et al. (2020)** and visualized in R (**RStudio Team 2019, R Core Team 2020**). The transcriptomic data and the array design were deposited in the Dryad repository (https://doi.org/10.5061/dryad.w3r2280xb).

## 5. ACKNOWLEDGEMENTS

The authors thank Yuji Yasukochi and Masanori Kobayashi for help with *Endoclita excrescens* karyotyping and rearing, Michal Rindoš and Michal Zapletal for moth collections, Dalibor Kodrík and Helena Štěrbová for providing specimens of *Pyrrhocoris aperus* for flow cytometry, Monika Kreklová and Magda Zrzavá for help with microscopy and Marie Korchová for assistance with rearing ghost moths. TJS wishes to thank Andrew Mitchell, Derek Smith and Dave Britton (Australian Museum, Sydney), You-Ning Su and Edward (Ted) Edwards (Australian National Insect Collection, Canberra), Cathy Burne (Tasmanian Museum and Art Gallery, Hobart), and Gary G. Anweiler (Strickland Museum, University of Alberta, Canada) for providing specimens for DNA extraction and sequencing; and Niklas Wahlberg, Eero J. Vesterinen, and Carlos Peña (then University of Turko), Alex Aitkin and Stephen Russel (Natural History Museum London) for help in the DNA lab, and Marie Djernæs (Aarhus University) for discussions and advice. We are also thankful to the Darwin Tree of Life project for genomic sequencing data from *Korscheltellus lupulina*. This project was funded by grants 20-20650Y (given to P.N.) and 17-13713S (given to F.M.) of the Czech Science Foundation, and a partially funded by a grant 23380030 from Japan Society for the Promotion of Science (given to K.S.). Financial support for DNA laboratory work (to T.J.S.) was provided by Australian Biological Resources Study (ABRS grant RF211-11). A.V. is thankful for the financial support to the Fulbright program (Fulbright-Masaryk grant 2016-28-14). Computational resources were provided by the “e-Infrastruktura CZ” project (e-INFRA LM2018140) under the Projects of Large Research, Development and Innovations Infrastructures program and the ELIXIR-CZ project (LM2018131), part of the international ELIXIR infrastructure.

## 6. DATA AVAILABILITY

The raw reads generated in this study have been deposited in the NCBI Sequence Read Archive (SRA) database under the Bioproject accession number PRJNA1012010. Other datasets generated in this study are available in the Dryad repository (https://doi.org/10.5061/dryad.w3r2280xb; reviewer sharing link https://datadryad.org/stash/share/7Hpd7meOmHavF0kiAs7VaTM_EY3JXSk5l55lcrQYCpg).

## SUPPLEMENTARY FIGURES

**Fig. S1.**
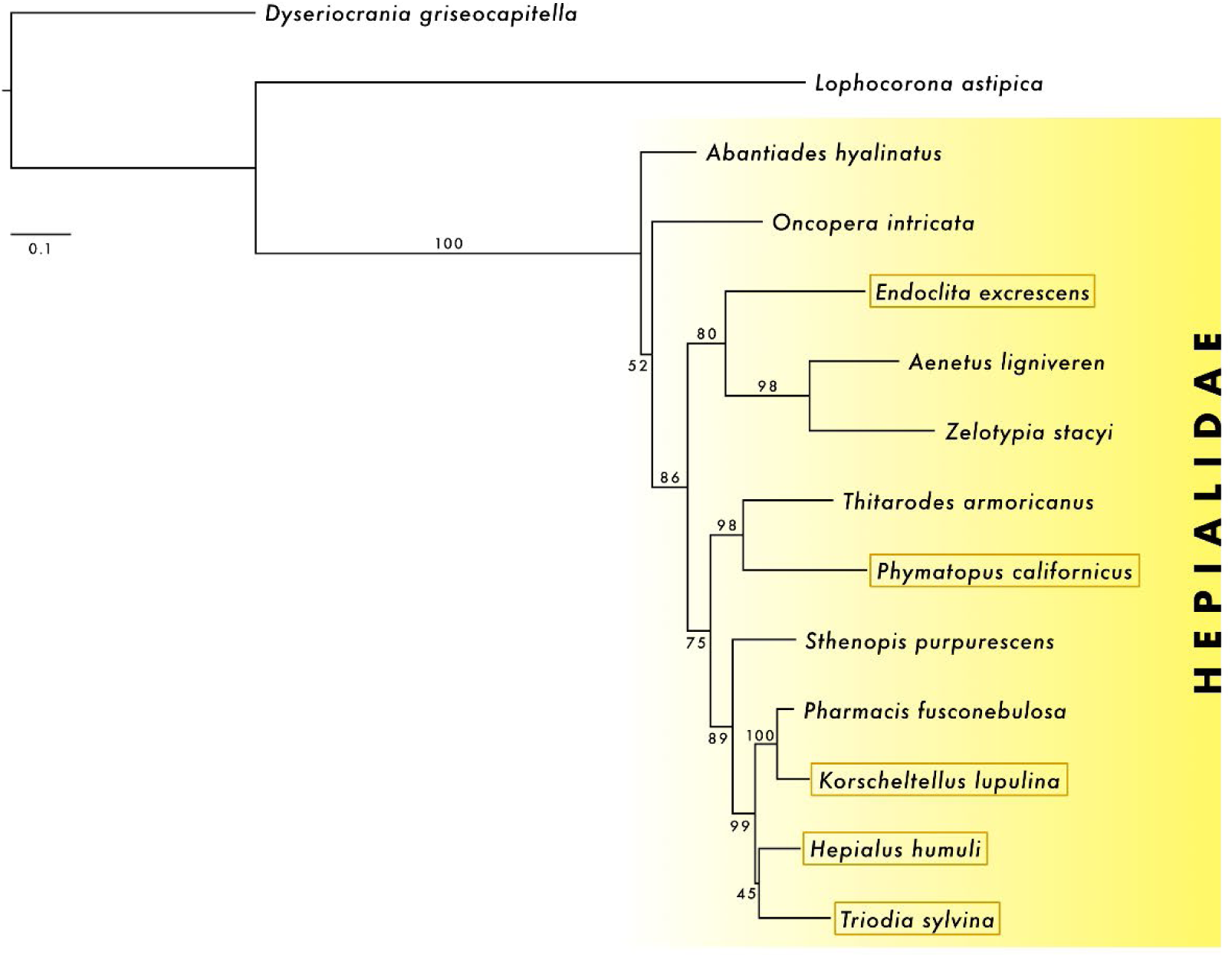
Phylogenetic tree of hepialid species resulting from Maximum likelihood analysis of one mitochondrial (*COI*) and six nuclear genes (*EF-1α, GAPDH, IDH, MDH, RpS5, wg*). The numbers at the nodes represent the bootstrap values. Species with a detected W chromosome are marked by a colored frame.

**Fig. S2.**
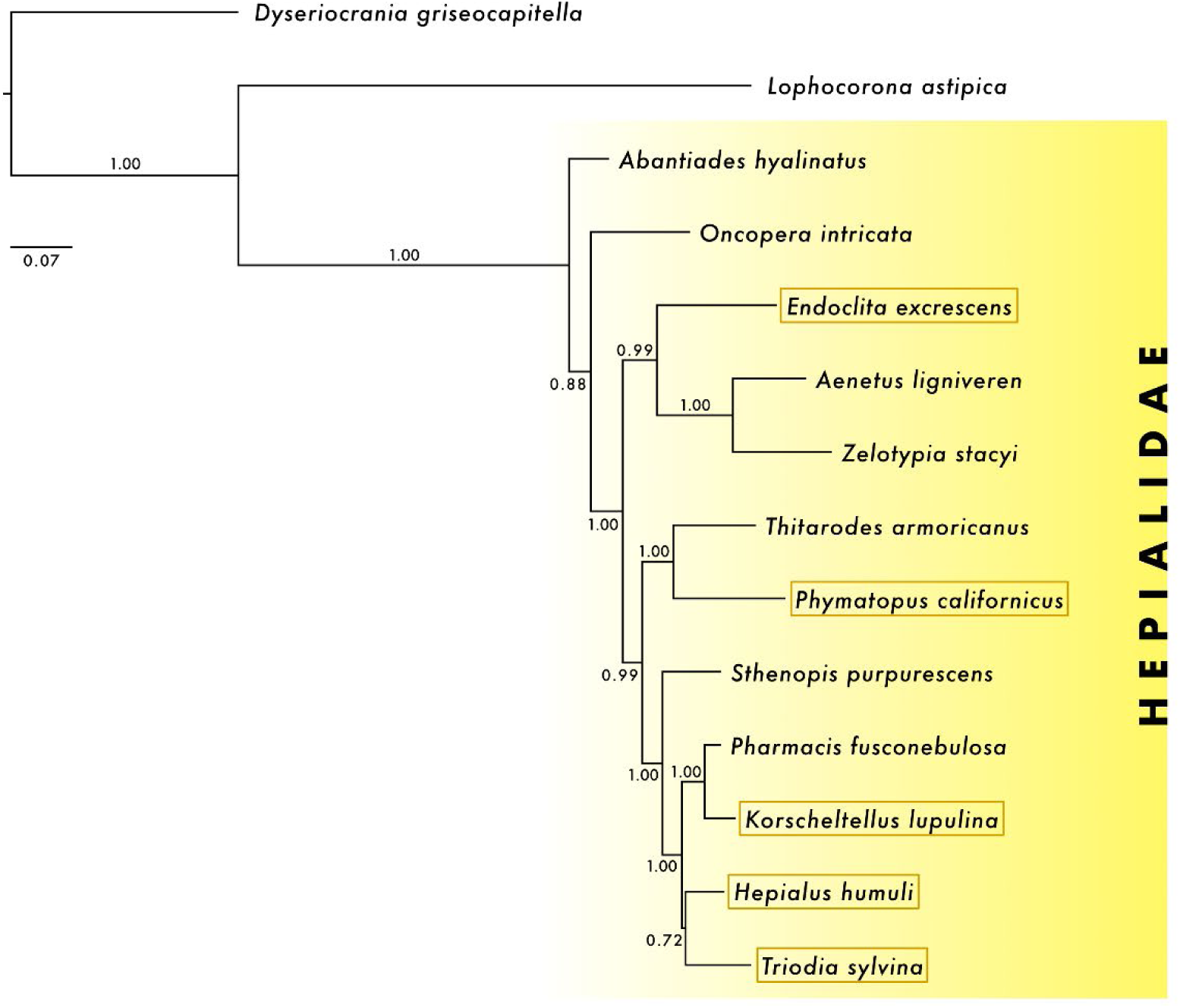
Phylogenetic tree of hepialid species resulting from Bayesian inference analysis of one mitochondrial (*COI*) and six nuclear genes (*EF1-α*, *GAPDH*, *IDH*, *MDH*, *RpS5*, *wg*). The numbers at the nodes represent the posterior probability values. Species with a detected W chromosome are marked by a colored frame.

**Fig. S3.**
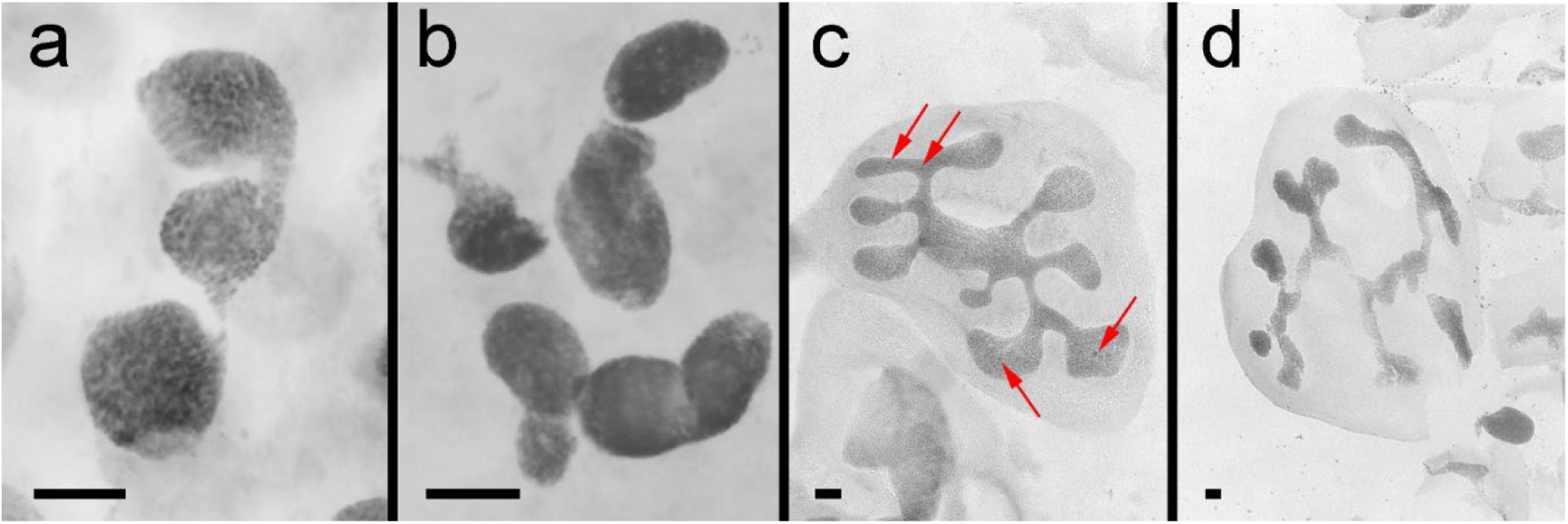
Sex chromatin assay in *Endoclita excrescens* (a–b) and *Phymatopus californicus* (c–d). Analysis of sex-specific heterochromatin in the polyploid nuclei of the Malpighian tubules stained with orcein revealed no heterochromatin bodies in females **(a,c)** or males **(b,d)** of either species. Examples of heterochromatin grains in female nuclei of *P. californicus* are marked by arrows. Bar = 20 µm.

**Fig. S4.**
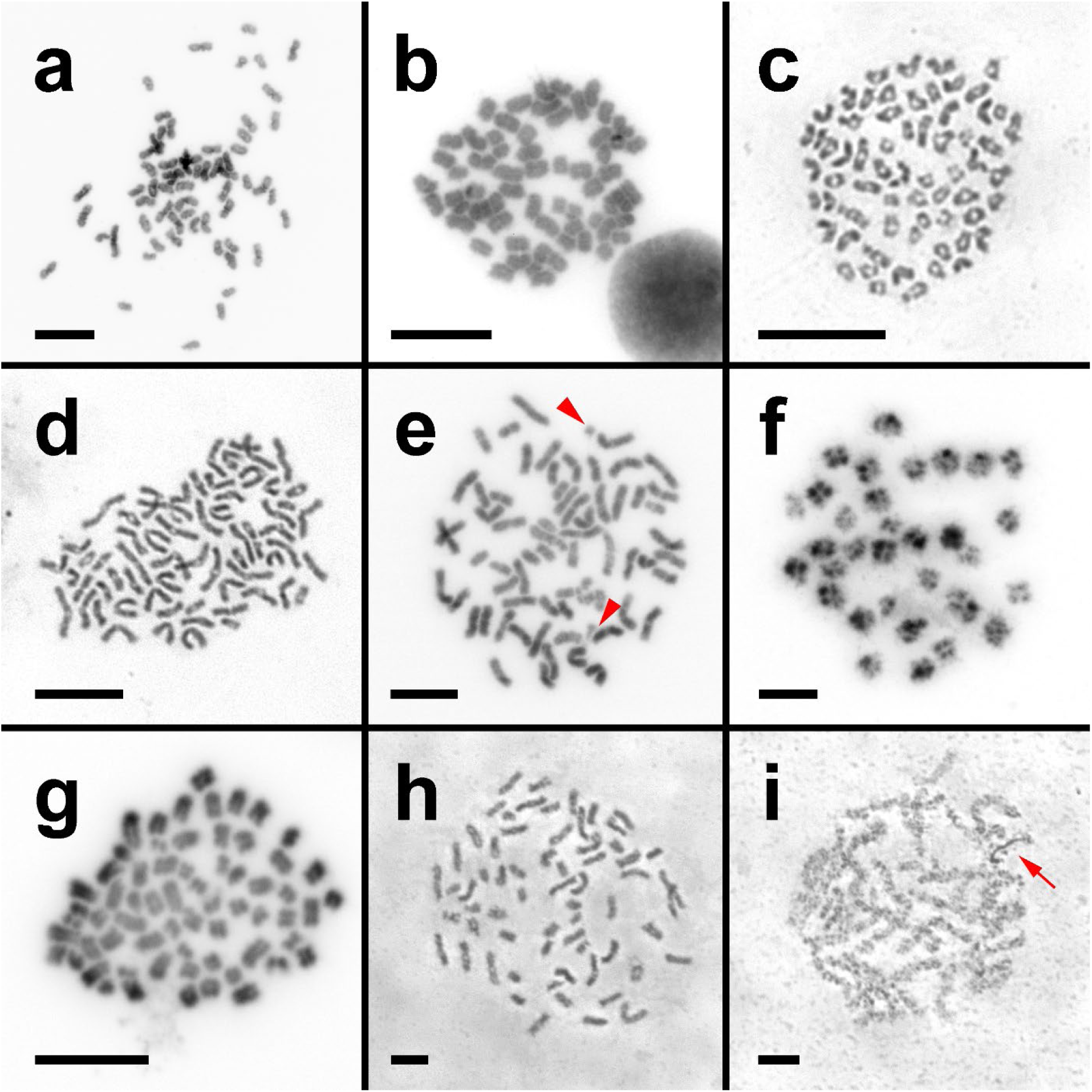
Mitotic and meiotic chromosomes of ghost moths. **(a, b)** Mitotic complements (2n = 64) of *Endoclita excrescens* – female metaphase **(a)** and male early anaphase **(b)**. **(c–e)** Mitotic complements (2n = 64) of *Hepialus humuli* – female early anaphase **(c)** and male prometaphases **(d,e)**. Note two supernumerary chromosomes (arrowheads) in **(e)**. **(f,g)** *Korscheltellus lupulina* – female nurse cell bivalents (n = 32) **(f)** and male mitotic early anaphase (2n = 64) **(g)**. **(h,i)** *Triodia sylvina* female – mitotic prometaphase (2n = 64) (h) and pachytene complement **(i)**. Note a deeply stained thread of the W chromosome (arrow) in **(i)**. Bar = 10 µm. Chromosomes in **(a,b, f,g)** stained with DAPI and images inverted, **(c–e, h,i)** with orcein.

**Fig. S5.**
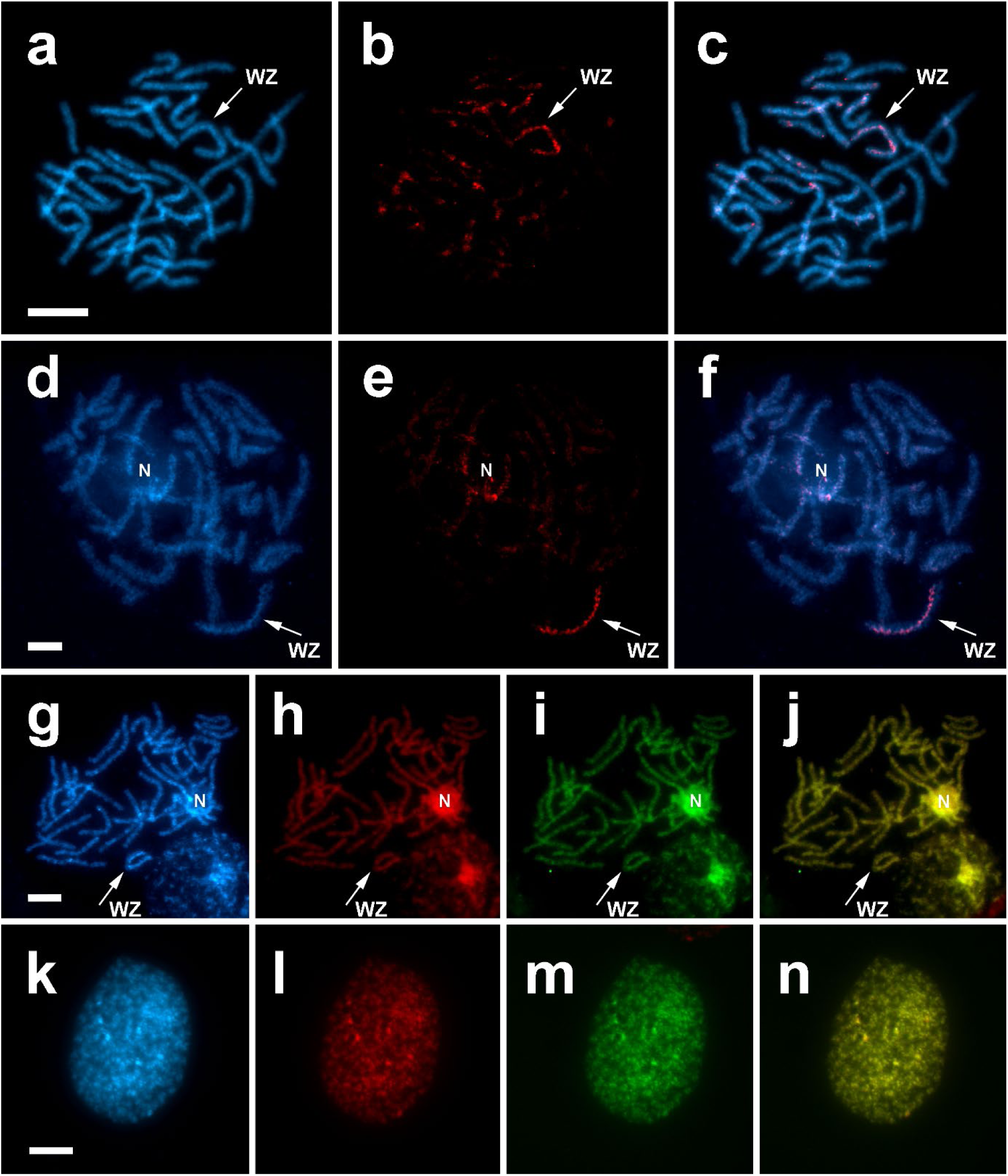
Detection of W chromosomes in pachytene oocytes of *Endoclita excrescens* (a–c), *Korscheltellus lupulina* (d–f), *Triodia sylvina* (g–j) and Hepialus humuli (k-n). Genomic *in situ* hybridization (GISH) revealed the presence of a differentiated W chromosome (red) in the WZ bivalent (arrow) of *E. excrescens* **(c)** and *K. lupulina* **(f)**. **(a,d)** Chromosomes stained with DAPI (blue); **(b,e)** hybridization signals of Cy3-labeled female gDNA probes (red); **(c, f)** merged images. With comparative genomic hybridization (CGH), the WZ bivalent in *Triodia sylvina* (arrow) was identified according to the DAPI-positive thread of the W chromosome **(g)**, but the W was not differentiated by Cy3-labeled female gDNA (**h**; red) and FITC-labeled male gDNA (**i**; green) probes. The merged image of both probes is shown in **(j)**. In *Hepialus humuli*, CGH have not revealed any differences between Cy3-labeled female gDNA (**l**; red) and FITC-labeled male gDNA (**m**; green) probes hybridized on female interphase nuclei, counterstained by DAPI (**k**; blue). The merged image of both probes is shown in **(n)**. N = nucleolus. Bar = 10 µm.

**Fig. S6.**
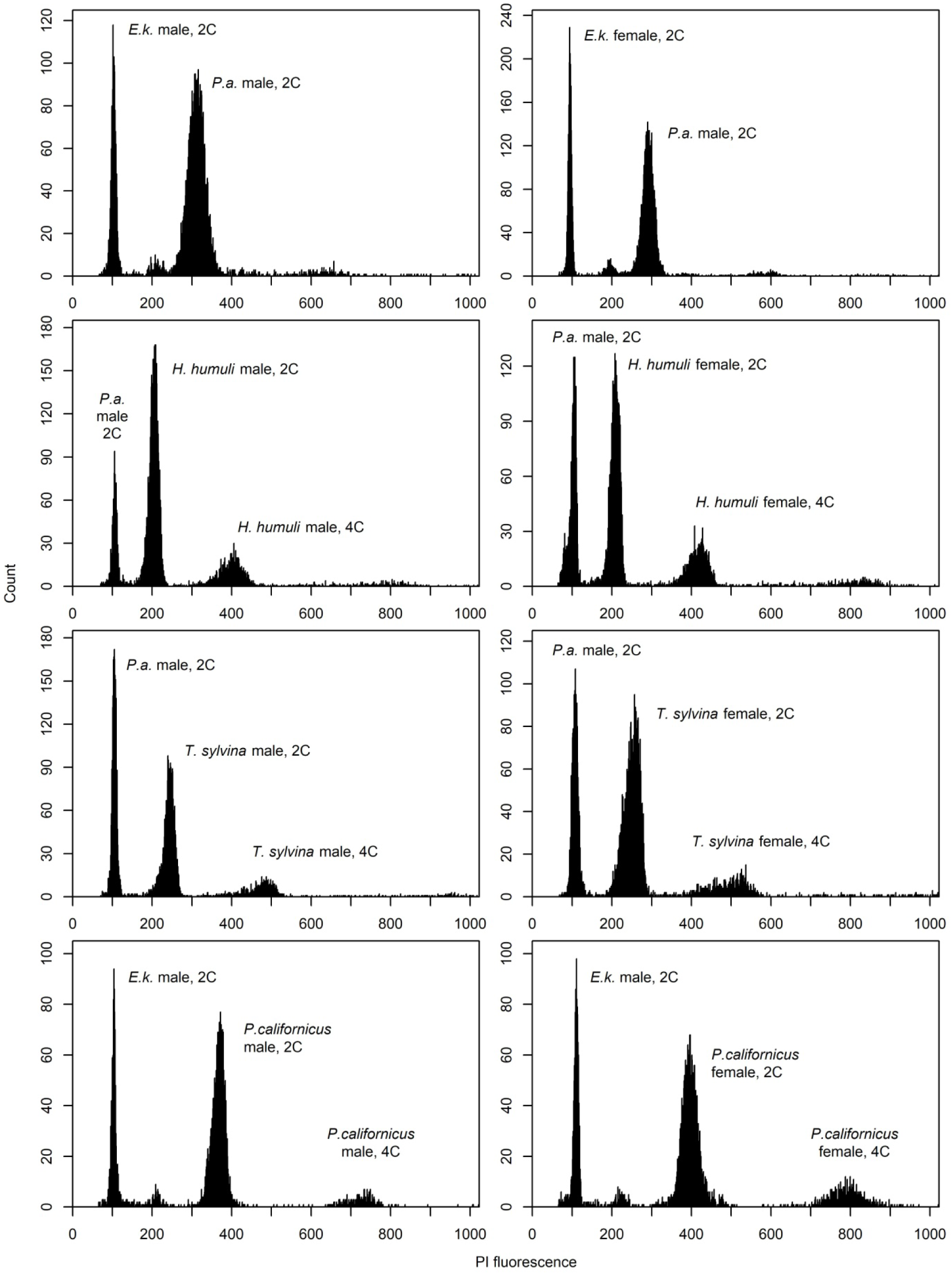
Examples of flow cytometric histograms in ghost moth species and the Mediterranean flour moth or the firebug, used as controls. Peaks of fluorescence corresponding to the internal standard and tested samples are clearly visible in all cases. *E.k.* – *Ephestia kuehniella*, *P.a.* – *Pyrrhocoris apterus*, *H. humuli* – *Hepialus humuli*, *T. sylvina* – *Triodia sylvina*, *P. californicus* – *Phymatopus californicus*. PI – propidium iodide.

## 10. Supplementary tables

**Table S1.**
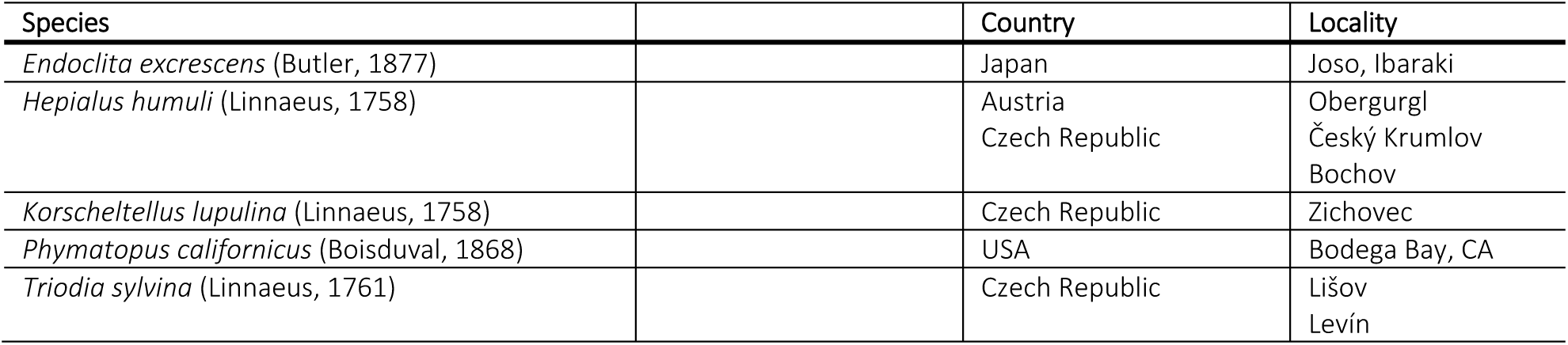
Origin of collected specimens of ghost moths.

**Table S2.**
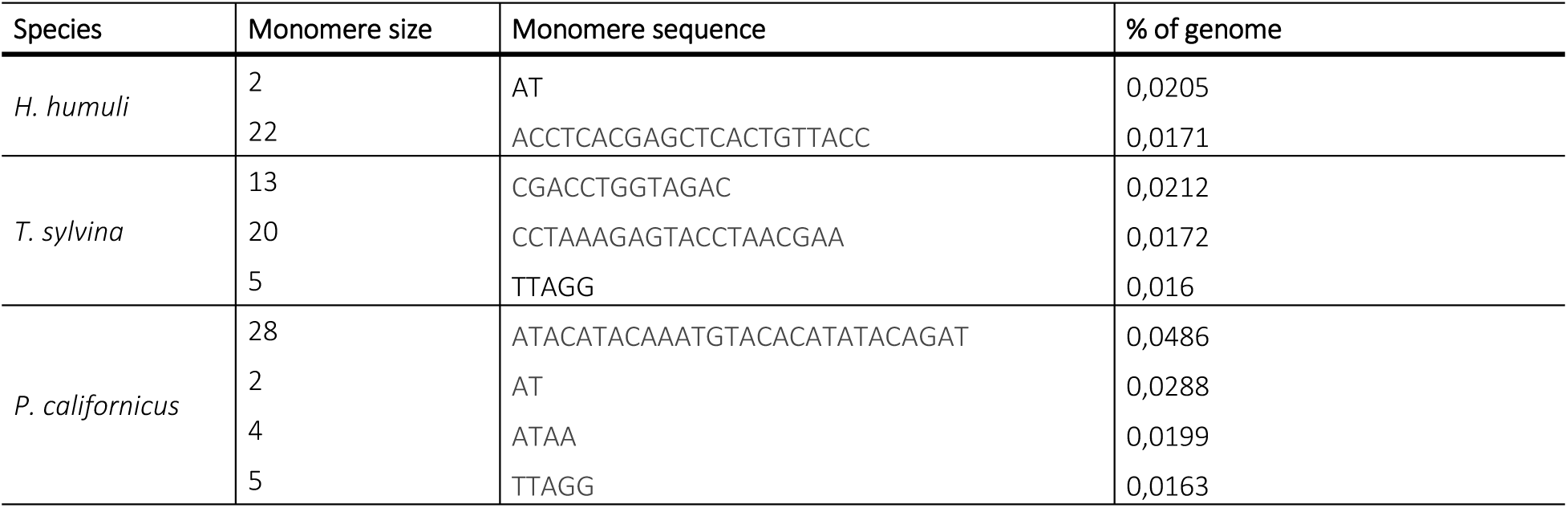
Micro and minisatellites in ghost moth species and their abundance.

**Table S3.**
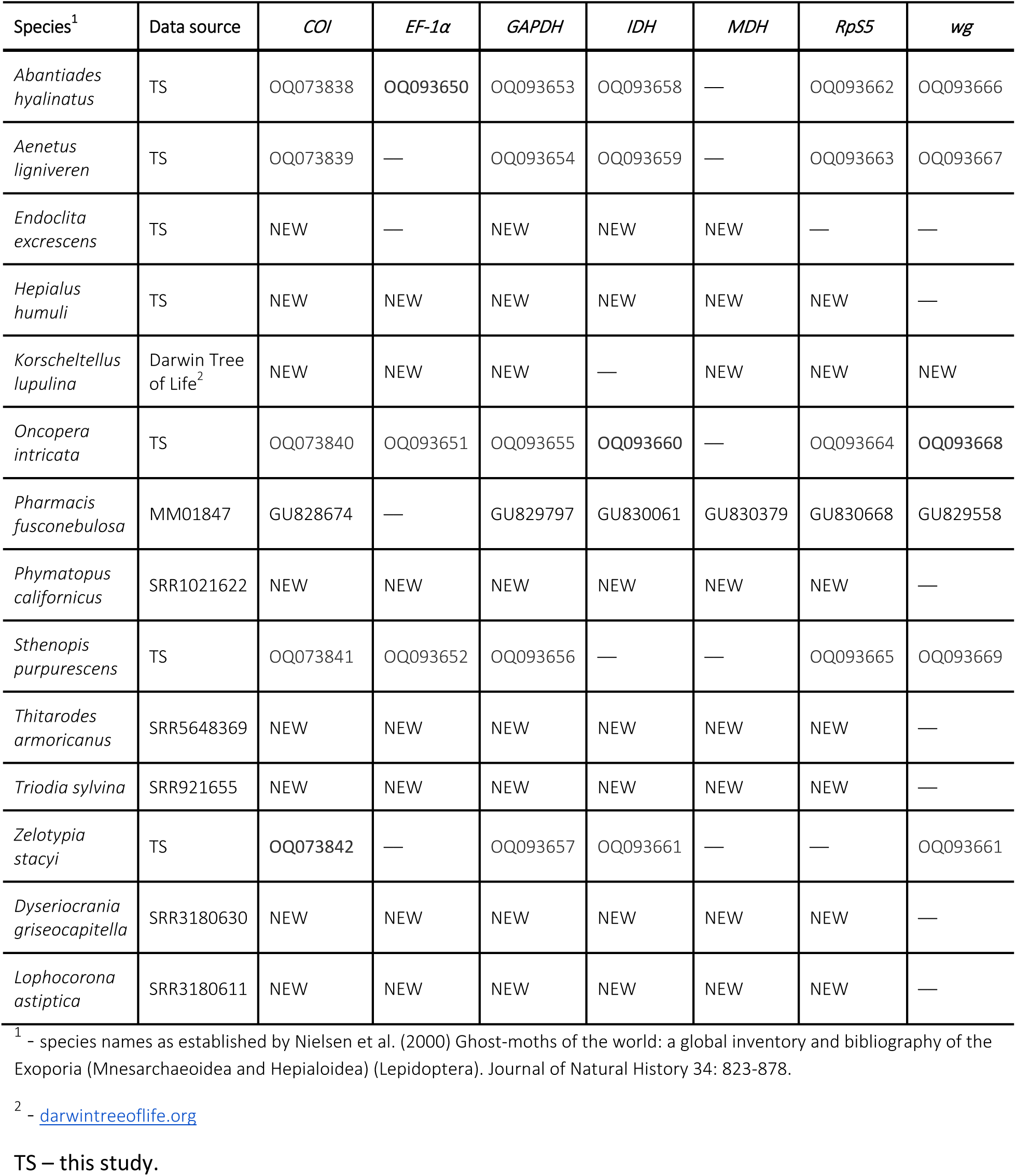
List of hepialid species, their IDs and GenBank accession numbers of gene markers used for the phylogenetic analysis.

**Table S4.** Partitioning schemes and models of molecular evolution selected for Maximum likelihood analysis (’models = raxml’) in PartitionFinder v. 2.1.1.

| Subset | Best Model | Partition names |
| --- | --- | --- |
| 1 | GTR+I+G | COI_pos1 |
| 2 | GTR+I+G | COI_pos2 |
| 3 | GTR+I+G | COI_pos3 |
| 4 | GTR+I+G | EF1a_pos1 |
| 5 | GTR+I+G | IDH_pos2, EF1a_pos2, GAPDH_pos2, RpS5_pos2, MDH_pos2 |
| 6 | GTR+G | EF1a_pos3 |
| 7 | GTR+G | GAPDH_pos1, wg_pos1, wg_pos2 |
| 8 | GTR+G | GAPDH_pos3 |
| 9 | GTR+G | IDH_pos1 |
| 10 | GTR+G | IDH_pos3 |
| 11 | GTR+G | MDH_pos1, RpS5_pos1 |
| 12 | GTR+G | MDH_pos3 |
| 13 | GTR+G | RpS5_pos3 |
| 14 | GTR+I+G | wg_pos3 |

**Table S5.**
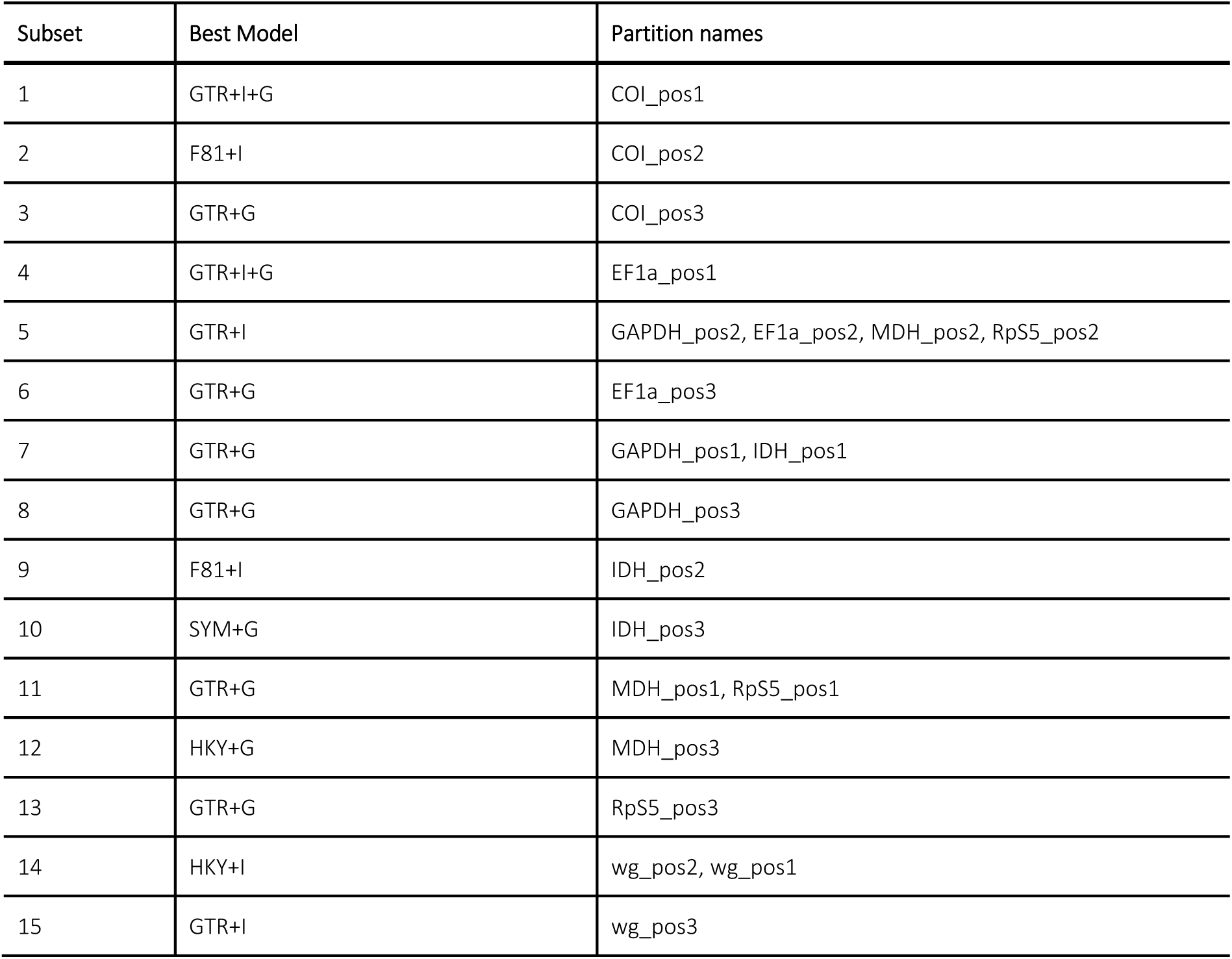
Partitioning schemes and models of molecular evolution selected for Bayesian analysis (’models = mrbayes’) in PartitionFinder v. 2.1.1.

| Subset | Best Model | Partition names |
| --- | --- | --- |
| 1 | GTR+I+G | COI_pos1 |
| 2 | F81+I | COI_pos2 |
| 3 | GTR+G | COI_pos3 |
| 4 | GTR+I+G | EF1a_pos1 |
| 5 | GTR+I | GAPDH_pos2, EF1a_pos2, MDH_pos2, RpS5_pos2 |
| 6 | GTR+G | EF1a_pos3 |
| 7 | GTR+G | GAPDH_pos1, IDH_pos1 |
| 8 | GTR+G | GAPDH_pos3 |
| 9 | F81+I | IDH_pos2 |
| 10 | SYM+G | IDH_pos3 |
| 11 | GTR+G | MDH_pos1, RpS5_pos1 |
| 12 | HKY+G | MDH_pos3 |
| 13 | GTR+G | RpS5_pos3 |
| 14 | HKY+I | wg_pos2, wg_pos1 |
| 15 | GTR+I | wg_pos3 |

## 11. SUPPLEMENTARY METHODS

### Data for molecular phylogeny

In order to obtain sequences necessary for reconstruction of phylogenetic relationships in Hepialidae *sensu stricto*, we used various sources of sequencing data, as well as amplification from genomic DNA.

#### 1. Transcriptomic data

*Hepialus humuli* transcriptome was sequenced de novo from the silk glands. Last instar *H. humuli* larvae were anaesthetized in water, and their silk glands were excised and stored at −80 °C. The frozen tissue was minced in liquid nitrogen, and total RNA was isolated using Trizol reagent (Life Technologies, Carlsbad, CA, USA). Subsequently, mRNA was enriched with Dynabeads Oligo(dT)25 (Ambion, Life Technologies). The rRNA was removed using the “A RiboMinus Eukaryote Kit for RNA-Seq” (Ambion, Austin, TX, USA). The cDNA library was prepared using the NEXTflex Rapid RNA-Seq Kit (Bioo Scientific, Austin, TX, USA). Sequencing was performed using MiSeq (Illumina, San Diego, USA), generating sequences in a 2 × 150 nt paired-end format. Data quality was checked using FastQC (Galaxy version 0.72 + galaxy1) and trimmed using Trimmomatic (Galaxy version 0.38.0). A total of 1.7 × 10^7^ reads were assembled de novo using Trinity software (version 2.9.1 + galaxy1, default settings) on the Galaxy platform (**Afgan et al., 2018**). The *H. humuli* transcriptome was deposited in the Dryad repository (https://doi.org/10.5061/dryad.w3r2280xb).

For *Phymatopus californicus*, *Thitarodes armoricanus*, *Triodia sylvina* and outgroups *Dyseriocrania griseocapitella* (Erinocraniidae) and *Lophocorona astiptica* (Lophocoronidae) we used transcriptomic data deposited in GenBank and assembled transcriptomes de novo. First, raw data were visualized in FastQC (v. 0.11.5; **Andrews 2010**) and low quality sequences and adaptor contamination was removed using Trimmomatic (v. 0.36; **Bolger et al. 2014**) with the following parameters: “ILLUMINACLIP: TruSeq3-PE-2.fa:2:30:10:1:true SLIDINGWINDOW:4:20 MINLEN:50”. Successful trimming was confirmed by FastQC and data were assembled de novo with Trinity (v. 2.6.5; **Grabherr et al. 2011**) with the deafult parameters.

Assembled transcriptomes were uploaded to GeneiousPrime (v. 2021.1.1; (https://www.geneious.com) and used to build a custom database for BLAST. Next, we searched GenBank for sequences of *Bombyx mori* genes *cytochrome oxidase subunit I* (*COI*; EU141360.1), *elongation factor 1 alpha* (*EF-1α*; EU136667.1), *glyceraldehyde-3-phosphate dehydrogenase* (*GAPDH*; EU141495.1), *isocitrate dehydrogenase* (*IDH*; EU141552.1), *cytosolic malate dehydrogenase* (*MDH*; EU141617.1), *ribosomal protein S5* (*RpS5*; EU141393.1) and *wingless* (wg; EU141241.1). These sequences were used as query for blastn to search the hepialid transcriptomes. In case the *B. mori* sequences yielded no hit, successfully retrieved sequences from other ghost moth species were used instead and the search was repeated.

#### 2. Expressed sequenced tags (ESTs)

In case of *Endoclita excrescens*, we used a library of expressed sequenced tags. The EST library construction and sequence were done according to **Kamimura et al (2012)** with a modification. The ESTs were searched using same approach as described in the previous section (1. Transcriptomic data).

#### 3. Genomic data

Sequences from *Korscheltellus lupulina* were obtained from whole genome sequencing data produced by the Darwin Tree of Life project (darwintreeoflife.org). Four datasets of 10X Linked-reads were downloaded from GitHub repository (https://github.com/darwintreeoflife/darwintreeoflife.data/blob/master/species/Korscheltellus_lupulina/Korscheltellus_lupulina.md), visualized in FastQC (v. 0.11.05; **Andrews 2010**) and filtered to remove low quality sequences and adaptor contamination with Trimmomatic (v 0.36; **Bolger et al. 2014**) using the following parameters and a custom file for adaptor removal (available at Dryad repository https://doi.org/10.5061/dryad.w3r2280xb): “ILLUMINACLIP:Truseq_PE_custom.fa:2:30:10:1:true SLIDINGWINDOW:5:20 CROP:135 HEADCROP:30 MINLEN:100”. Quality of the filtered data was verified with FastQC. Next, the reads were aligned in an unpaired manner to sequences of target genes from *P. californicus* (see above) and *Pharmacis fusconebulosa* (GenBank accession number GU829558) with bowtie2 and “--very-sensitive-local” settings. Alignments of mapping reads were used to reconstruct the marker sequences in GeneiousPrime (v. 2021.1.1; (https://www.geneious.com).

#### 4. Amplification from genomic DNA

Material for DNA work was provided by Andrew Mitchell, Derek Smith and Dave Britton (Australian Museum, Sydney), You-Ning Su and Ted Edwards (Australian National Insect Collection, Canberra), Cathy Burne (Tasmanian Museum and Art Gallery, Hobart), and Gary G. Anweiler (Strickland Museum, University of Alberta, Canada). Genomic DNA was extracted from one or two legs of *Aenetus ligniveren*, *Abantiades hyalinatus*, *Zelotypia stacyi*, *Oncopera intricata* and *Sthenopis purpurescens* using DNeasy Bood & Tissue kit (Qiagen, Hilden, Germany) according to the manufacturer’s instruction with the following exceptions: the lysis was performed for approximately 1 8h at 38°C (**Krosch & Cranston 2012**), and elution was done twice, using 100 µl elution buffer for each step.

PCR reactions targeting the genes *COI*, *EF-1α*, *GAPDH*, *IDH*, *RpS5* and *wingless* were performed with the general profile: initial denaturation for 7 min at 96 °C; 40 cycles of 30 sec denaturation at 96°C; 30 sec annealing at 45-55°C; 2 min extension at 72°C, followed by a final extension period of 10 min at 72°C. Annealing temperature was 45°C for COI, 50°C for *wingless*, and 55°C for the remaining genes. For majority of markers, the universal-tail primers from **Wahlberg & Wheat (2008)** were used. Due to amplification problems in some specimen, new internal primers (without universal tails) were designed for *GAPDH* and *IDH*. All primers are listed in the table below (Table SM1).

**Table SM1.**
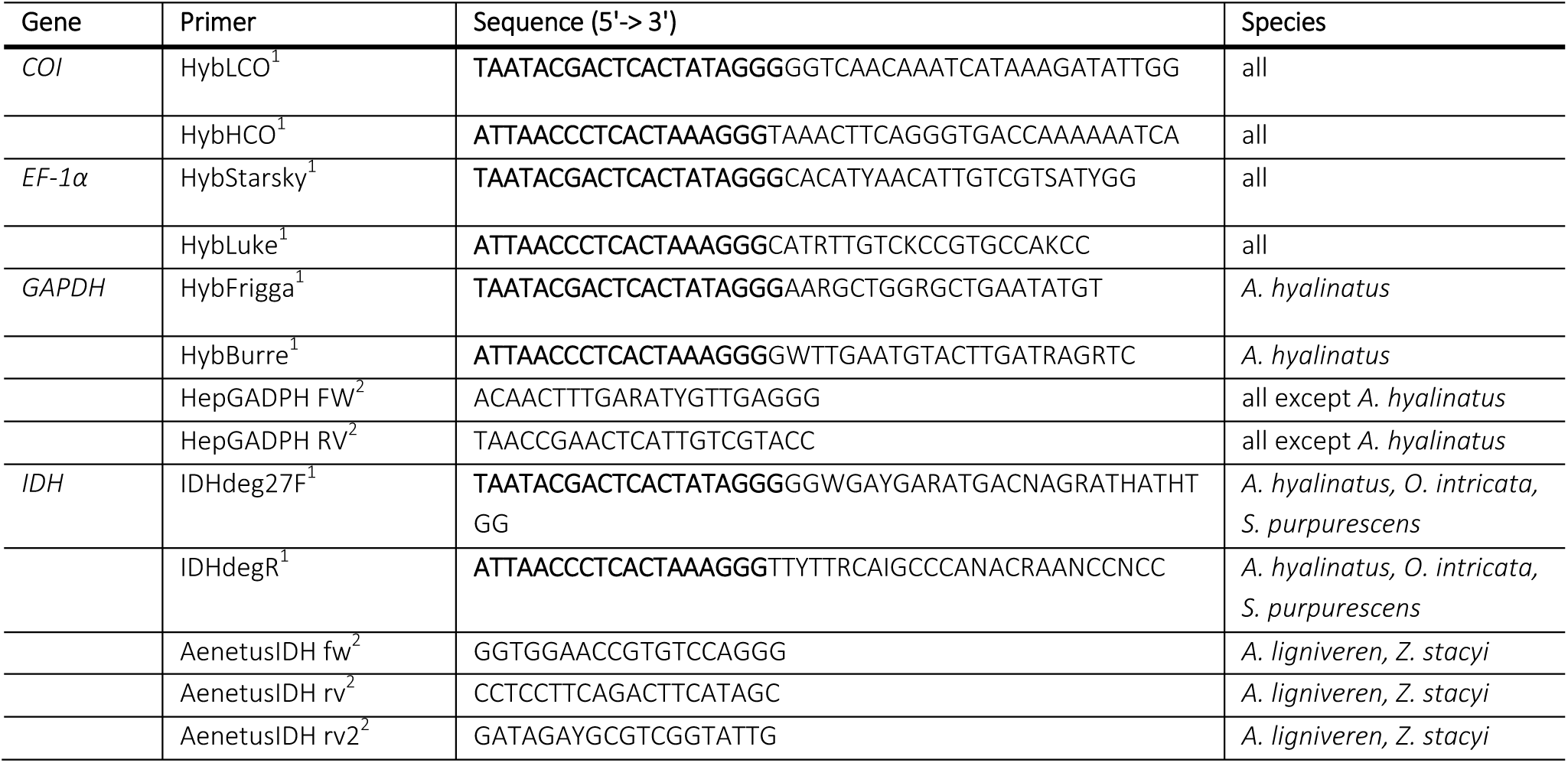
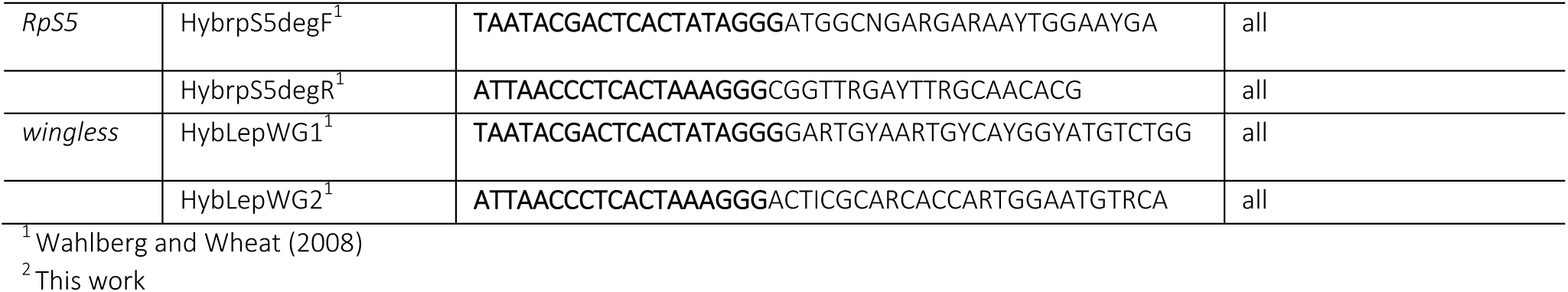
Primers used for amplification of phylogenetic markers. Universal tails indicated in bold.

Direct bidirectional Sanger sequencing with BigDye (Applied Biosystems, Waltham, MA, USA) was done using either the universal primer tails (primers from **Wahlberg & Wheat 2008**) or PCR primers (internal *GAPDH* and *IDH* primers). Consensus sequences for each gene were constructed in Lasergene SeqMan 7.0 (DNASTAR, Madison, WI, USA) based on bidirectional sequence reads.

## 12. SUPPLEMENTARY DATA

### Supplementary data 1

**Partial sequences of *Phymatopus californicus cytochrome C oxidase subunit I* (*COI*) gene, corresponding to the ancestral (anc) and the neo-W chromosome (neo) chromosomal races.** Note three single nucleotide polymorphisms indicated in the alignment.

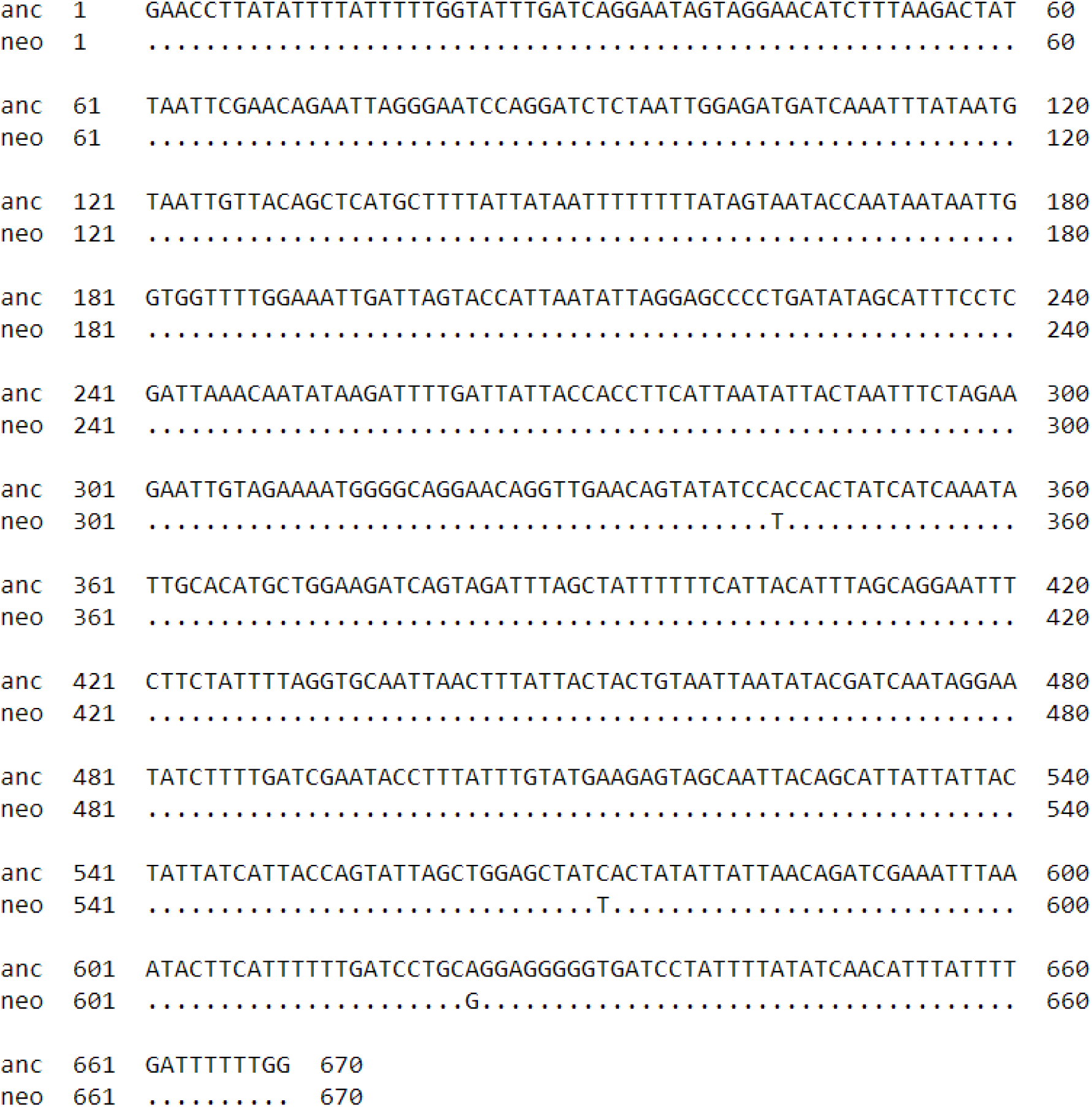

